# A Tad-like apparatus is required for contact-dependent prey killing in predatory social bacteria

**DOI:** 10.1101/2021.02.25.432843

**Authors:** Sofiene Seef, Julien Herrou, Paul de Boissier, Laetitia My, Gael Brasseur, Donovan Robert, Rikesh Jain, Romain Mercier, Eric Cascales, Bianca Habermann, Tâm Mignot

## Abstract

*Myxococcus xanthus*, a soil bacterium, predates collectively using motility to invade prey colonies. Prey lysis is mostly thought to rely on secreted factors, cocktails of antibiotics and enzymes, and direct contac with *Myxococcus* cells. In this study, we show that on surfaces the coupling of A-motility and contact-dependent killing is the central predatory mechanism driving effective prey colony invasion and consumption. At the molecular level, contact-dependent killing involves a newly discovered type IV filament-like machinery (Kil) that both promotes motility arrest and prey cell plasmolysis. In this process, Kil proteins assemble at the predator-prey contact site, suggesting that they allow tight contact with prey cells for their intoxication. Kil-like systems form a new class of Tad-like machineries in predatory bacteria, suggesting a conserved function in predator-prey interactions. This study further reveals a novel cell-cell interaction function for bacterial pili-like assemblages.

## Introduction

Bacterial predators have evolved strategies to consume other microbes as a nutrient source. Despite the suspected importance of predation on microbial ecology^1^, a limited number of bacterial species are currently reported as predatory. Amongst them, obligate intracellular predators collectively known as BALOs *(eg Bdellovibrio bacteriovorus*)^1^, penetrate their bacterial prey cell wall and multiply in the periplasm, escaping and killing the host bacteria^2^. Quite differently, facultative predators (meaning that they can be cultured in absence of prey if nutrient media are provided, *ie Myxococcus, Lysobacter* and *Herpetosiphon*^1^) attack their preys extracellularly, presumably by secreting antimicrobial substances and digesting the resulting products. Among these organisms and studied here, *Myxococcus xanthus*, a delta-proteobacterium, is of particular interest because it uses large-scale collective movements to attack prey bacteria in a so-called “wolf-pack” mechanism^3^.

A tremendous body of work describes how *Myxococcus* cells move and respond to signals in pure culture^4^. In contrast, mechanistic studies of the predatory cycle have been limited. Currently, it is considered that coordinated group movements allow *Myxococcus* cells to invade prey colonies and consume them via the secretion of a number of diffusible factors, extracellular enzymes, antibiotics and outer membrane vesicles^3,5,6^. While each of these processes could contribute to predation, evidence for their requirement is still missing^3^. In addition, *Myxococcus* cells have also been observed to induce prey cell plasmolysis upon contact^7^. While a number of contact-dependent mechanisms could be involved, including Type VI secretion^8^ and Outer Membrane Exchange (OME^9^, see below), none have yet been implicated in predation. In this study, we analyzed the importance of motility and contact-dependent killing in the *Myxococcus* predation cycle.

To explore these central questions, we first developed a sufficiently resolved imaging assay where the *Myxococcus* predation cycle can be imaged stably at the single cell level over periods of time encompassing several hours with a temporal resolution of seconds. The exact methodology underlying this technique is described in a dedicated manuscript^10^; briefly, the system relates predatory patterns observed at the mesoscale with single cell resolution, obtained by zooming in and out on the same microscopy specimen (Figure 1). Here, we employed it to study how *Myxococcus* cells invade and grow over *Escherichia coli* prey cells during the initial invasion stage (Figure 1, Movie S1).

**Figure 1.**
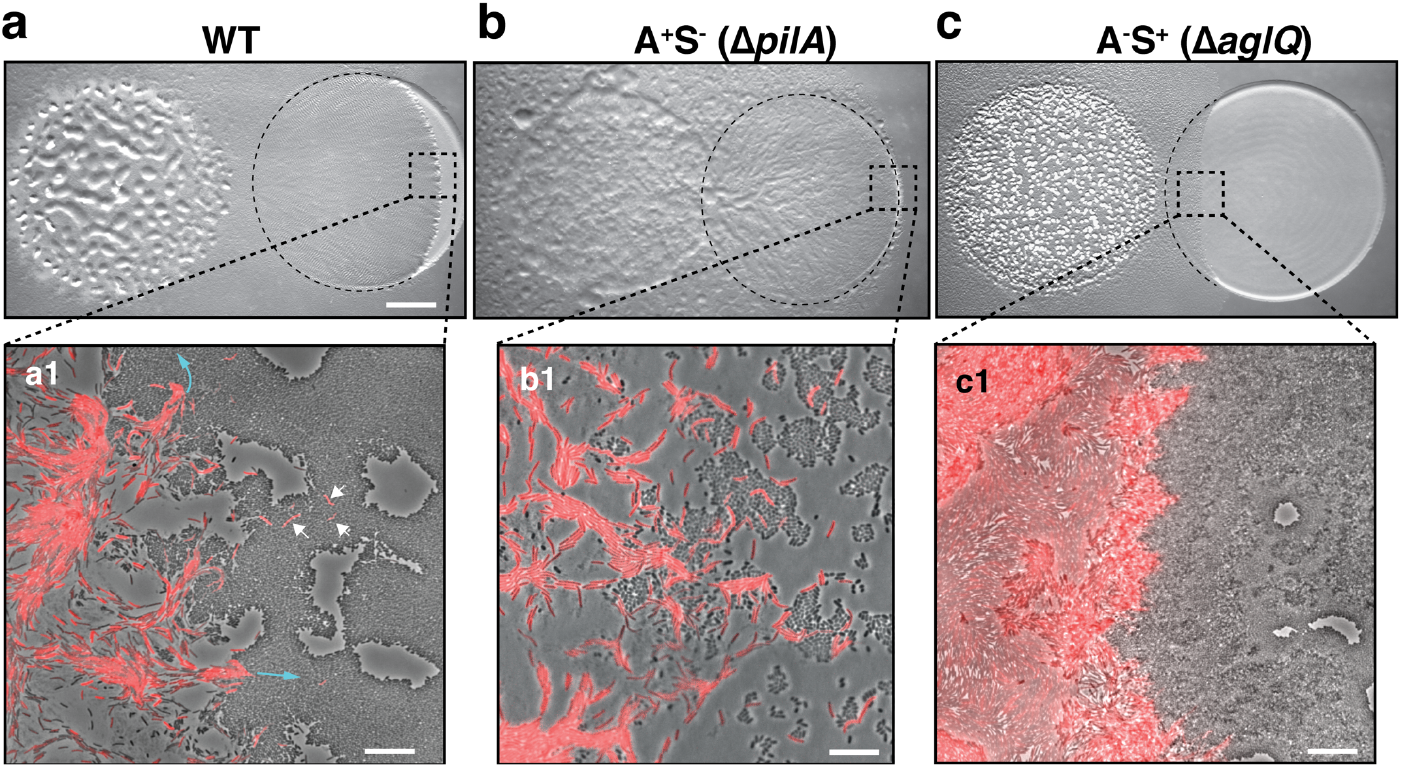
A-motility is required for invasion of prey colonies. Colony plate assays showing invasion of an *E. coli* prey colony (dotted line) 48 hours after plating by WT **(a, Movie S1)**, A^+^S^-^ **(b)** and A^-^S^+^ **(c, Movie S2)** strains. Scale bar = 2 mm. **a1:** Zoom of the invasion front. *Myxococcus* single cells are labelled with mCherry. Blue arrows show the movement of “arrowhead” cell groups as they invade prey colonies. White arrows point to A-motile single cells that penetrated the prey colony. Scale bar = 10 µm. See associated Movie S1 for the full time lapse. **b1:** Zoom of the invasion front formed by A^+^S^-^ cells. The A-motile *Myxococcus* cells can infiltrate the prey colony and kill prey cells. Scale bar = 10 µm. **c1:** Zoom of the invasion front formed by A^-^S^+^ cells. Note that the S-motile *Myxococcus* cells come in contact with the prey colony, but in absence of A-motility, the predatory cells fail to infiltrate the colony and remain stuck at the border. Scale bar = 10 µm. See associated Movie S2 for the full time lapse.

### A-motility is required for prey colony invasion

Although the function of motility in prey invasion is generally accepted, *Myxococcus xanthus* possesses two independent motility systems and the relative contribution of each system to the invasion process is unknown. Social (S)-motility is a form of bacterial “twitching” motility that uses so-called Type IV pili (TFP) acting at the bacterial pole^11^. In this process, polymerized TFPs act like “grappling hooks” that retract and pull the cell forward. S-motility promotes the coordinated movements of *Myxococcus* cells within large cell groups due to interaction with a self-secreted extracellular matrix formed of Exo-Polysaccharide (EPS)^12–14^. A(Adventurous)-motility promotes the movement of *Myxococcus* single cells at the colony edges. A-motility is driven by a mobile cell-envelope motor complex (named Agl-Glt) that traffics in helical trajectories along the cell axis, driving rotational propulsion of the cell when it becomes tethered to the underlying surface at so-called bacterial Focal Adhesions (bFAs)^15^. We tested the relative contribution of each motility system to prey invasion by comparing the relative predatory performances of WT, A^+^S^-^(Δ*pilA*^16^) and A^-^(Δ*aglQ*^16^) S^+^ (Figure 1). Interestingly, although A^+^S^-^ cells were defective in the late developmental steps (fruiting body formation), they were still proficient at prey invasion (Figure 1b). On the contrary, the A^-^S^+^ strain was very defective at prey colony invasion (Figure 1c). Zooming at the prey colony border, it was apparent that the A^-^S^+^ cells were able to expand and contact the prey colony, but they were unable to penetrate it efficiently, suggesting that Type IV pili on their own are not sufficient for invasion (Figure 1c, Movie S2). Conversely, A-motile cells were observed to penetrate the tightly-knitted *E. coli* colony with single *Myxococcus* cells moving into the prey colony, followed by larger cell groups (Figure 1a). Similar motility requirements were also observed in a predatory assay where predatory and prey cells are pre-mixed before they are spotted on an agar surface (Figure S1a). Thus, A-motility is the main driver of prey invasion on surfaces.

### *Invading* Myxococcus *cells kill prey cells upon contact*

To further determine how A-motile cells invade the prey colony, we shot single cell time-lapse movies of the invasion process. First, we localized a bFA marker, the AglZ protein^17^ fused to Neon-Green (AglZ-NG) in *Myxococcus* cells as they penetrate the prey colony. AglZ-NG binds to the cytoplasmic face of the Agl-Glt complex and has long been used as a bFA localization marker; it generally forms fixed fluorescent clusters on the ventral side of the cell that retain fixed positions in gliding cells^17^. As *Myxococcus* cells invaded prey colonies, they often formed “arrow-shaped” cell groups, in which the cells within the arrow assembled focal adhesions (Figure 2a, Movie S3). *E. coli* cells lysed in the vicinity of the *Myxococcus* cells, suggesting that a contact-dependent killing mechanism as reported by Zhang *et al*.^7^ occurs during prey colony invasion (Figure 2a). To observe this activity directly, we set up a *Myxococcus* - *E. coli* interaction microscopy assay where predator - prey interactions can be easily studied, isolated from a larger multicellular context (see Methods). In this system, A-motile *Myxococcus* cells were observed to mark a pause and disassemble bFAs when contacting *E. coli* cells (Figure 2b, Movie S4, further quantified below); this pause was invariably followed by the rapid death of *E. coli*, as detected by the instantaneous dispersal of a cytosolic fluorescent protein (mCherry or GFP, Figure 2b-2c, observed in n=20 cells). This observation suggest that the killing occurs by plasmolysis, a process which is likely to be the same as that described by Zhang *et al*.^7^. To demonstrate this, we mixed *Myxococcus* cells with *E. coli* cells in which peptidoglycan (PG) had been labeled by fluorescent D-amino Acids (TADA^15^). TADA is covalently incorporated into the PG pentapeptide backbone and it does not diffuse laterally^18^. We first observed contraction of the *E. coli* cytosolic dense region at the pole by phase contrast (Figure 2d), which was followed by the appearance of a dark area in the PG TADA staining exactly at the predator-prey contact site (Figure 2d). It is unlikely that this dark area forms due to the new incorporation of unlabeled prey PG, because it was detected immediately upon prey cell death and propagated bi-directionally afterwards (Figure 2d-e). Thus, these observations suggest that, upon contact, *Myxococcus* induces degradation of the *E. coli* PG, which provokes cell lysis due to loss of turgor pressure and hyper osmotic shock^7^. The bi-directional propagation of PG hydrolysis (as detected by loss of TADA signal) suggests that PG hydrolysis could be driven by the activity of PG hydrolase(s) disseminating from the predator-prey contact site.

**Figure 2.**
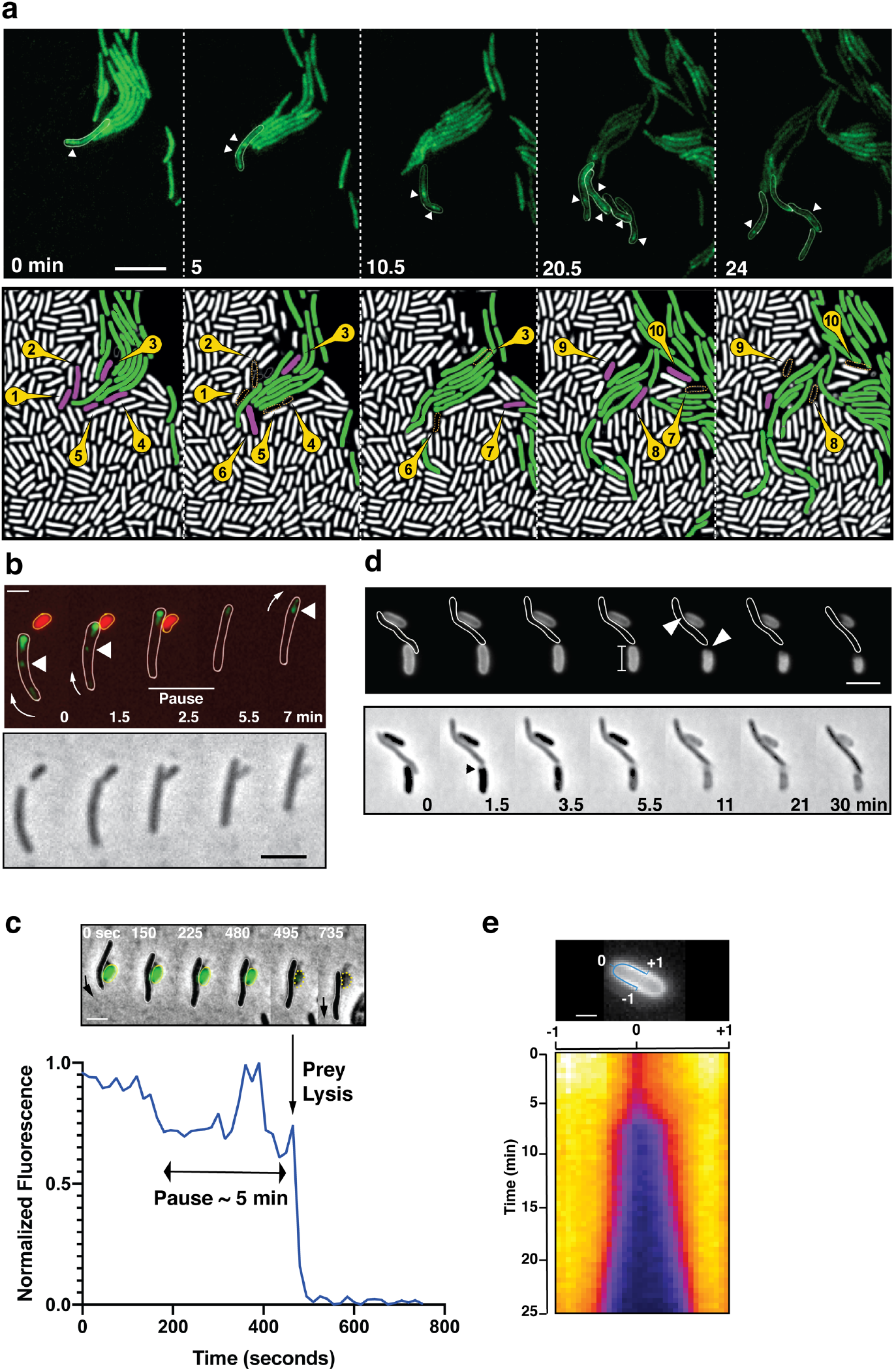
A-motile cells kill prey cells by contact. **a:** Prey (*E. coli*) colony invasion by an “arrowhead formation”. Activity of the A-motility complex is followed by monitoring *Myxococcus* cells expressing the bFA-localized AglZ-YFP protein. Upper panel: Cells within the arrowhead (examples shown in white) assemble bFAs (white arrowheads). Lower panel: Semantic segmentation (see methods) of the total cell population, *E. coli* (white) and *Myxococcus* (green). The numbered and colored *E. coli* cells (magenta) are the ones that are observed to lyse as the Myxococcus cells penetrate the colony. See associated Movie S3 for the full time lapse. Scale bar = 10 µm. **b:** bFAs are disassembled when *Myxococcus* establishes lytic contacts with prey cells. Shown is an AglZ-YFP expressing *Myxococcus* cell establishing contact with an mCherry-expressing *E. coli* cell (overlay and phase contrast image). Note that the *Myxococcus* cell resumes movement and thus re-initiates bFA formation immediately after *E. coli* cell lysis. See associated Movie S4 for the full time lapse. Scale bar = 2 µm. **c:** *Myxococcus* (outlined in white) provoke *E. coli* plasmolysis. Top: shown is a GFP-expressing *E. coli* cell lysing in contact with a *Myxococcus* cell. GFP fluorescence remains stable for 5 min after contact and becomes undetectable instantaneously, suggesting plasmolysis of the *E. coli* cell. Scale bar = 2 µm. Bottom: graphic representation of fluorescence intensity loss upon prey lysis. **d**,**e :** *Myxococcus* contact provoke local degradation of the *E. coli* peptidoglycan. **d:** *E. coli* PG was labeled covalently with the fluorescent D-amino acid TADA. Two *E. coli* cells lyse upon contact. Holes in the PG-labelling are observed at the contact sites (white arrows). Note that evidence for plasmolysis and local IM membrane contraction is visible by phase contrast for the lower *E. coli* cell (dark arrow). Scale bar = 2 µm. **e:** Kymograph of TADA-labeling corresponding to the upper *E. coli* cell. At time 0 which corresponds to the detection of cell lysis, a hole is detected at the contact site and propagates bi-directionally from the initial site showing that the prey cell wall is degraded in time after cell death. Scale bar = 1 µm.

### A predicted Tad-pilus is required for contact-dependent killing

We next aimed to identify the molecular system that underlies contact-dependent killing. Although motility appears to be essential during the predation process (Figure S1a), at the microscopic level, direct transplantation of A^-^S^-^ (*aglQ pilA*) in *E. coli* prey colonies still exhibit contact-dependent killing (Figure S1a-b), demonstrating that the killing activity is not carried by the motility complexes themselves. *Myxococcus xanthus* also expresses a functional Type VI secretion system (T6SS), which appears to act as a factor modulating population homeostasis and mediating Kin discrimination between *M. xanthus* strains^8,19^. A T6SS deletion strain (Δ*t6ss*) had no observable defect in contact-dependent killing of prey cells (Figure S1a and S1c). In addition, the *Myxococcus* T6SS assembled in a prey-independent manner as observed using a functional VipA-GFP strain that marks the T6SS contractile sheath^20^ (Figure S1d-f), confirming that T6SS is not involved in predatory killing on surfaces.

To identify the genes (directly or indirectly) involved in the contact-dependent killing mechanism, we designed an assay where contact-dependent killing can be directly monitored in liquid cultures and observed via a simple colorimetric assay. In this system, the lysis of *E. coli* cells can be directly monitored when intracellular β-galactosidase is released in buffer containing ChloroPhenol Red-β-D-Galactopyranoside (CPRG), which acts as a substrate for the enzyme and generates a dark red hydrolysis reaction product^21^. Indeed, while *Myxococcus* or *E. coli* cells incubated alone did not produce color during a 120-hour incubation, their mixing produced red color indicative of *E. coli* lysis after 24 h (Figure S2a). In this assay, a *t6SS* mutant was still able to lyse *E. coli* cells, demonstrating that it does not report on T6SS-dependent killing (Figure S2a-b). CPRG hydrolysis was not detected when *Myxococcus* and *E. coli* were separated by a semi-permeable membrane that allows diffusion of soluble molecules, showing that the assay reports contact-dependent killing (Figure S2a). In this liquid assay, the *Myxococcus* - *E. coli* contacts are very distinct from contacts on solid surfaces and thus, the genetic requirements are likely quite distinct. Indeed, in the liquid CPRG assay, we observed that TFPs are essential for killing while the Agl/Glt system is dispensable (Figure S2b). In this condition, TFPs promote a prey-induced aggregation of cells (Figure S2c) and thus probably mediate the necessary tight contacts between *Myxococcus* and *E. coli* cells. As shown below, the killing process itself is the same in liquid as the one observed on surfaces and it is not directly mediated by the TFPs.

Given the probable indirect effect of TFPs, we next searched additional systems involved in CPRG contact-dependent killing, using a targeted approach and testing the effect of mutations in genome annotated cell-envelope complexes, on contact-dependent killing in liquid cultures. Doing so, we identified two critical genetic regions, the MXAN_3102-3108 and the MXAN_4648-4661 (Figure 3). Functional annotations indicate that both genetic regions carry a complementary set of genes encoding proteins that assemble a so-called **T**ight **ad**herence (Tad) pilus. Bacterial Tad pili are members of the type IV filament superfamily (also including Type IV pili, a and b types, and Type II secretion systems) and extrude polymeric pilin filaments assembled via inner membrane associated motors through an OM secretin^22^. Tad pili have been generally involved in bacterial adhesion and more recently, in contact-dependent regulation of adhesion^23^. Within the MXAN_3102-3108 cluster, genes with annotated functions encode a predicted pre-pilin peptidase (CpaA and renamed KilA) following the *Caulobacter crescentus* Tad pilus encoding *cpa* genes nomenclature), a secretin homolog (CpaC/KilC) and a cytoplasmic hexameric ATPase (CpaF/KilF) (Figure 3a, Figure S3a-b, Table S1). All the other genes encode proteins of unknown function, with two predicted OM lipoproteins and several proteins containing predicted ForkHead-Associated domains (FHA^24^, Table S1, see discussion). The second genetic region, MXAN_4648-4661, contains up to 14 predicted open-reading frames of which the only functionally annotated genes encode homologs of the Tad IM platform proteins (CpaG/KilG and CpaH/KilH), OM protein (CpaB/KilB), major pilin (Flp/KilK) and two pseudo-pilin subunits (KilL, M) (Figure 3a, Figure S3c-d, Table S1). However, the splitting of Tad homologs in distinct genetic clusters is a unique situation^22^ and asks whether these genes encode proteins involved in the same function.

**Figure 3.**
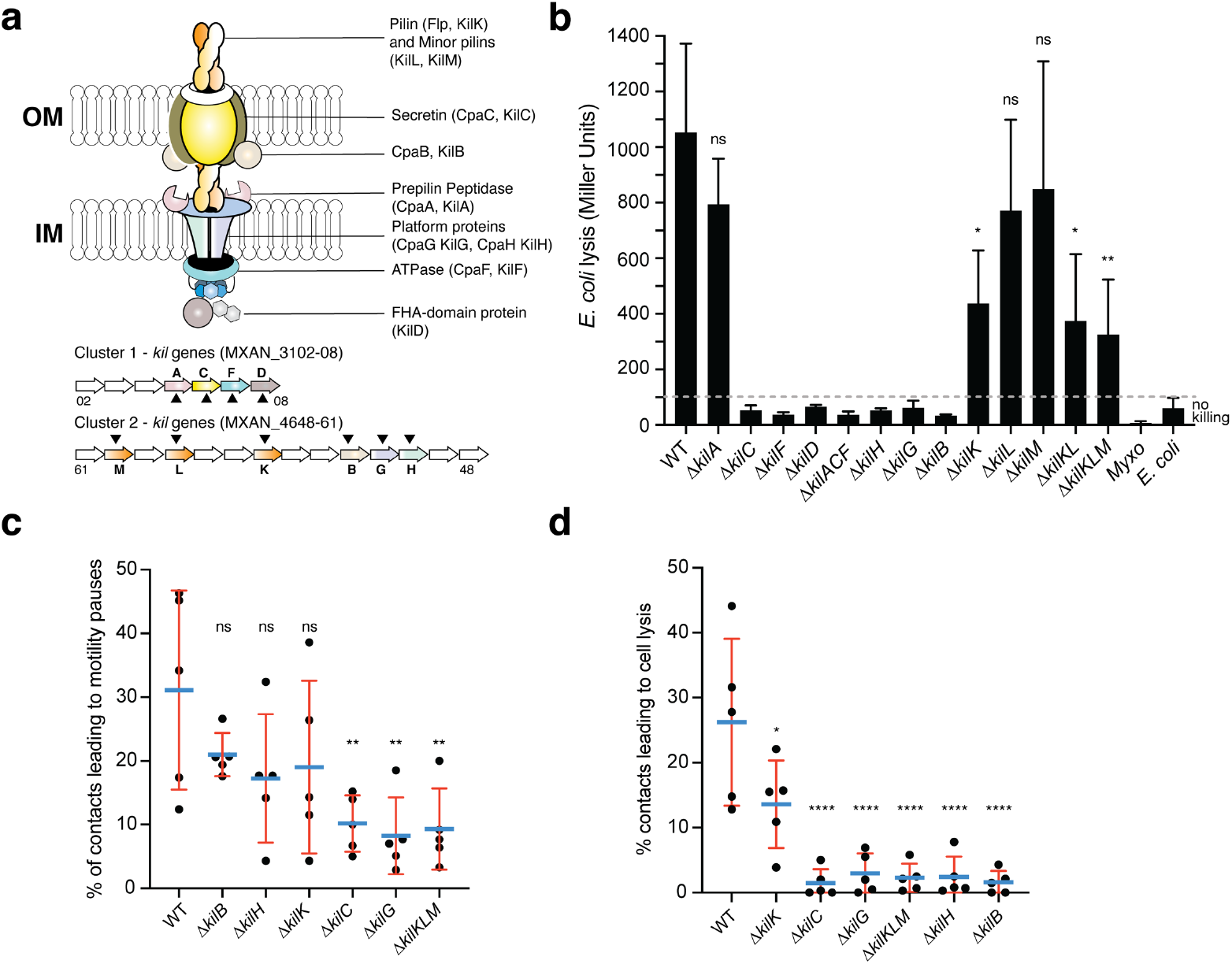
A Tad-like apparatus is required for prey recognition and contact-dependent killing. **a:** Model structure of the Kil system following bioinformatics predictions. Annotated cluster 1 and cluster 2 genes are shown together with the possible localization of their protein product. Dark triangles indicate the genes that were deleted in this study. **b:** *kil* mutants are impaired in *E. coli* lysis in liquid. Kinetics CPRG-hydrolysis by ß-Galactosidase (expressed as Miller Units) observed after co-incubation of *Myxococcus* WT and *kil* mutants and *E. coli* for 24 hours. *M. xanthus* and *E. coli* alone were used as negative controls. This experiment was performed independently four times. **c:** The percentage of contacts with *E. coli* leading to a pause in motility was calculated for *M. xanthus* wild-type (from five independent predation movies, number of contacts observed n= 807) and *kil* mutants (number of contacts observed in Δ*kilC*: n= 1780; Δ*kilH*: n= 1219; Δ*kilG*: n=1141; Δ*kilB*: n= 842; Δ*kilK*: n=710; Δ*kilKLM*: n= 1446) **d:** The percentage of contacts with *E. coli* leading to cell lysis was also estimated. In panels (b), (c) and (d), error bars represent the standard deviation of the mean. One-way ANOVA statistical analysis followed by Dunnett’s posttest was performed to evaluate if the differences observed, relative to wild-type, were significant (*: p≤0.05, **: p≤0.01, ****: p≤0.0001) or not (ns: p>0.05).

Expression analysis suggests that the cluster 1 and cluster 2 genes are expressed together and induced in starvation conditions (Figure S4a). We systematically deleted all the predicted Tad components in cluster 1 and 2 alone or in combination and measured the ability of each mutant to lyse *E. coli* in the CPRG colorimetric assay (Figure 3b). All the predicted core genes, IM platform, OM secretin and associated CpaB homolog are essential for prey lysis, with the exception of the putative pre-pilin peptidase, KilA. Deletion of the genes encoding predicted pseudo-pilins KilL and M did not affect *E. coli* killing; in these conditions, pilin fibers are only partially required because deletion of KilK, the major pilin subunit, reduces the lytic activity significantly but not fully (Figure 3b). Given that the genes are organized into potential operon structures, we confirmed that the CPRG-killing phenotypes of predicted cluster 1 and cluster 2 core genes were not caused by potential polar effects (Figure S4b). In liquid, predicted core gene mutants had the same propensity as wild-type to form biofilms in presence of the prey suggesting that they act downstream in the interaction process (Figure S2c).

We next tested whether liquid killing and contact-dependent killing on surfaces reflected the same process. For this, we analyzed selected *kil* mutants, predicted secretin (KilC), IM platform (KilH and KilG), OM-CpaB homolog (KilB), pilin and pseudopilins (KilK, L, M) in contact-dependent killing at the single cell level. Prey recognition is first revealed by the induction of a motility pause upon prey cell contact (Figure 2). This recognition was severely impaired although not fully in secretin (*kilC*), IM platform protein (*kilG*) and triple pilin (Δ*kilKMN*) mutants (Figure 3c, ∼8% of the contacts led to motility pauses vs ∼30% for the WT). In contrast, recognition was not impaired to significant levels in IM platform protein (*kilH*), CpaB-homolog (*kilB*) and pilin (*kilK*) mutants (Figure 3c). The potential basis of this differential impact is further analyzed in the discussion. On the contrary, prey cell plasmolysis was dramatically impacted in all predicted core components (∼2% of the contacts led to prey lysis vs ∼26% for the WT), the only exception being the single pilin (*kilK*) mutant in which prey cell lysis was reduced but still present (∼13%, Figure 3d). Deletion of all three genes encoding pilin-like proteins nevertheless affected in prey cell killing to levels observed in core component mutants. This is not observed to such extent in the CPRG assay, which could be explained by different cell-cell interaction requirements, perhaps compensation by TFPs in liquid cultures. Given the prominent role of the pilins at the single cell level, the predicted pre-pilin peptidase KilA would have been expected to be essential. However, expression of the *kilA* gene is very low under all tested conditions (Figure S4a). Prepilin peptidases are known to be promiscuous^25^ and thus another peptidase (ie PilD, the Type IV pilus peptidase^26^) could also process the Kil-associated pilins. This hypothesis could however not be tested because PilD appears essential for reasons that remain to be determined^26^. Altogether, the data supports that the proteins from the two clusters function in starvation conditions and that they could make up a Tad-like core structure, for prey cell recognition, regulating motility in contact with prey cells, and prey killing, allowing contact-dependent plasmolysis.

### Kil proteins assemble at contact sites and mediate motility regulation and killing

We next determined if the Kil proteins indeed form a single Tad-like system in contact with prey cells. To do so, the predicted ATPAse (KilF) (Figure 3a) was N-terminally fused to the Neon Green (NG) and expressed from the native chromosomal locus. The corresponding fusion appeared fully functional (Figure S4c). In absence of prey cells, NG-KilF was diffuse in the cytoplasm. Remarkably, when *Myxoccocus* cells established contact with prey cells, NG-KilF rapidly formed a fluorescent-bright cluster at the prey contact site. Cluster formation was invariably followed by a motility pause and cell lysis. Observed clusters did not localize to any specific cellular site but they formed where *Myxococcus* cells touched prey cells. Cluster formation was correlated to motility arrest and their dispersal coincided with motility resumption (Figure 4a, Movie S5). To confirm that the KilF clusters reflect assembly of a full Tad-like apparatus, we next attempted to label a component of the IM platform KilG (Cluster 2), expressing a KilG-NG fusion this time ectopically from a *pilA* promoter (*PpilA*) in a *kilG* mutant background. The fusion was partially functional (Figure S4d), but nevertheless KilG-NG clusters could also be observed forming at the prey contact site immediately after lysis (Figure 4b, Movie S6). These results strongly suggest that a Tad-apparatus assembled from the products of the cluster 1 and cluster 2 genes.

**Figure 4.**
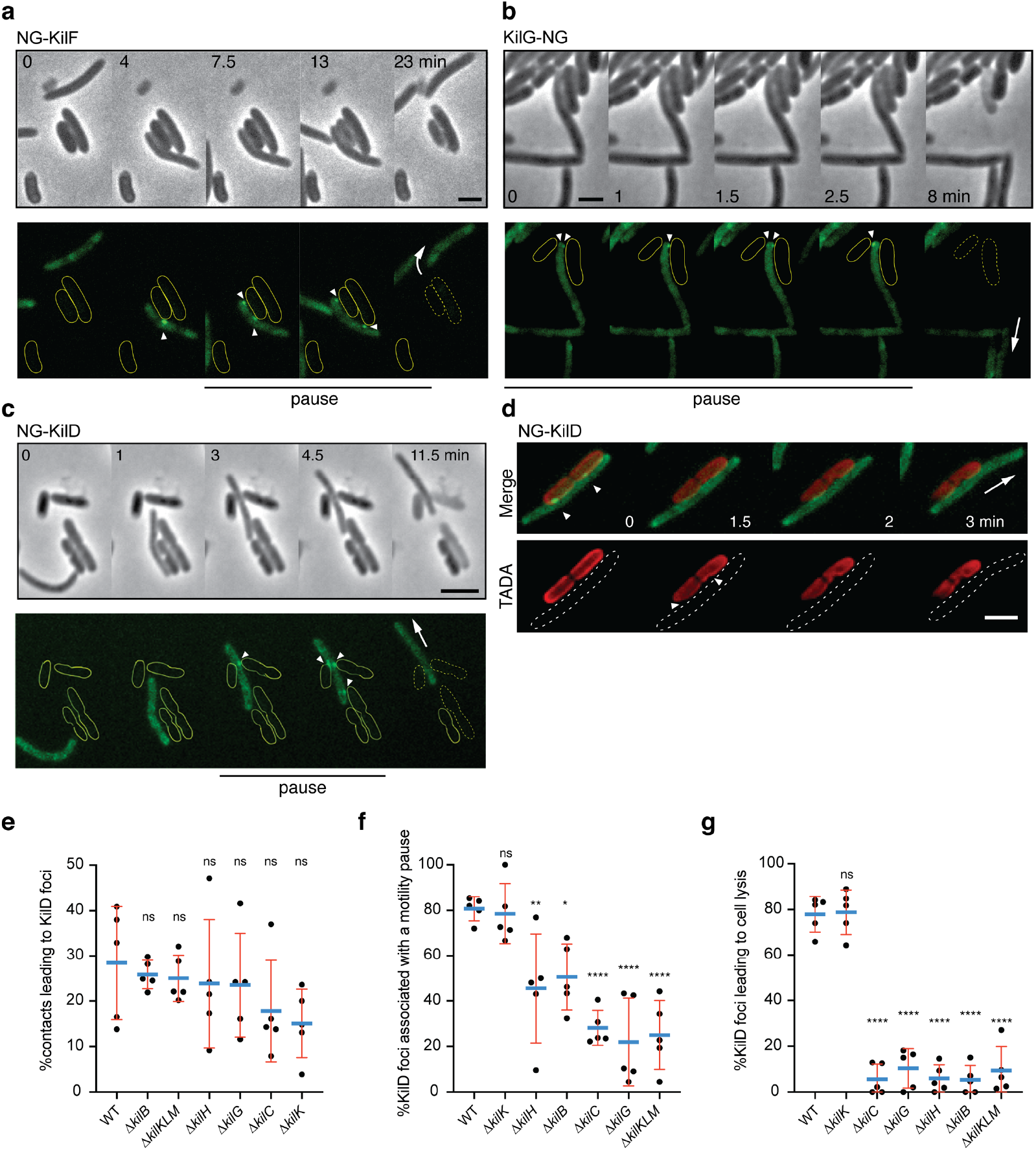
The Kil Tad-like system assembles upon contact and causes prey cell lysis. **a:** NG-KilF clusters form in contact with the prey and their formation precedes cell lysis. Scale bar = 2 µm. See associated Movie S5 for the full time lapse. **b:** KilG-NG forms clusters at the contact site with the prey and their formation is followed by the prey cell lysis. Scale bar = 2 µm. See associated Movie S6 for the full time lapse. **c:** NG-KilD clusters only form in contact with the prey and their formation precedes cell lysis. Scale bar = 2 µm. See associated Movie S7 for the full time lapse. **d:** PG-holes are formed at the cluster-assembly sites. Representative picture of TADA-labelled *E. coli cells* in the presence of NG-KilD expressing *Myxococcus xanthus* cells. PG holes and clusters are indicated with white arrows. Scale bar = 2 µm. **e**: The percentage of contacts with *E. coli* leading to KilD foci formation was calculated for *M. xanthus* wild-type (from five independent predation movies, number of contacts observed n= 807) and *kil* mutants (number of contacts observed in Δ*kilC*: n= 1780; Δ*kilH*: n= 1219; Δ*kilG*: n=1141; Δ*kilB*: n= 842; Δ*kilK*: n=710; Δ*kilKLM*: n= 1446). **f**: The percentage of KilD foci associated with a motility pause was also estimated for *M. xanthus* WT (from five independent predation movies, number of NG-KilD foci observed n= 198) and *kil* mutants (number of NG-KilD foci observed in Δ*kilC*: n= 320; Δ*kilH*: n= 270; Δ*kilG*: n= 251; Δ*kilB*: n= 215; Δ*kilK*: n= 94; Δ*kilKLM*: n= 355). **g**: The percentage of KilD foci leading to *E. coli* lysis was estimated as well. In panels (d), (e) and (f), error bars represent the standard deviation to the mean. One-way ANOVA statistical analysis followed by Dunnett’s posttest was performed to evaluate if the differences observed, relative to wild-type, were significant (*: p≤0.05, **: p≤0.01, ****: p≤0.0001) or not (ns: p>0.05).

Additional non-core proteins are also recruited at the contact sites: downstream from *kilF* and likely co-transcribed, the MXAN_3108 gene (*kilD*, Figures 3a and S4a) encodes a predicted cytoplasmic multidomain protein also required for killing and thus functionally associated with the Kil apparatus (Figure 3b). An NG-KilD fusion was fully functional, also forming a fluorescent-bright cluster at a prey contact site, followed by motility arrest and prey cell lysis (Figure 4c and S4c, Movie S7). Given that this protein is the most downstream component of the cluster 1 region, which facilitates further genetic manipulations (see below), we next used it as a reporter for Kil system-associated functions for further characterization and in-depth quantifications. First, to confirm that prey intoxication occurs at sites where the Kil proteins are recruited, we imaged NG-KilD in the presence of *E. coli* cells labeled with TADA. As expected, PG degradation was detection at the points where the clusters are formed, showing that cluster formation correlates with contact dependent killing (Figure 4d). Using cluster assembly as a proxy for activation of the Kil system, we measured that killing is observed within ∼2 min after assembly, a rapid effect which suggests that Kil system assembly is tightly connected to a prey cell lytic activity (Figure S4e).

We next used NG-KilD as a proxy to monitor the function of the Kil Tad apparatus in prey recognition and killing. For this, NG-KilD was stably expressed from the native chromosomal locus in different genetic backgrounds (Figure S4f). In WT cells, NG-KilD clusters only formed in the presence of prey cells and ∼30% contacts were productive for cluster formation (Figure 4e). In *kil* mutants, NG-KilD clusters still formed upon prey cell contact with a minor reduction (up to ∼2 fold in the KilC and KilK), suggesting theTad-like apparatus is not directly responsible for initial prey cell sensing (Figure 4e, Movie S8). Nevertheless, cluster assembly was highly correlated to motility pauses (Figure 4f); which was impaired (up to 60%) in the *kil* mutants (except in the pilin, *kilK* mutant) and most strongly in the *kilC* (secretin), *kilG* (IM platform) and triple pilin (*kilKLM)* mutants (Figure 4f). Strikingly and contrarily to WT cells, cluster formation was not followed by cell lysis in all *kil* mutants, except in the major pilin *kilK* mutant (or very rarely, ∼4% of the time versus more than 80% in WT, Figure 4g). Altogether, these results indicate the Tad-like Kil system is dispensable for immediate prey recognition, but functions downstream to induce a motility pause and critically, provoke prey cell lysis.

### The Kil apparatus is central for Myxococcus predation

We next tested the exact contribution of the *kil* genes to predation and prey consumption. This question is especially relevant because a number of mechanisms have been proposed to contribute to *Myxococcus* predation and all involve the extracellular secretion of toxic cargos^9,11,12^. In pure cultures, deletion of the *kil* genes is not linked to detectable motility and growth phenotypes, suggesting that the Tad-like Kil system mostly operates in predatory context (Figure S4g-h). Critically, core *kil* mutants where unable to predate colonies on plate, which could be fully complemented when corresponding *kil* genes were expressed ectopically (Figure 5a). When observed by time lapse, a *kil* mutant (here Δ*kilACF*) can invade a prey colony, but no prey killing is observed, showing that the prey killing phenotype is indeed due to the loss of contact-dependent killing (Figure 5b, Movie S9). To measure the impact of this defect quantitatively, we developed a FACS-based assay that directly measures the relative proportion of *Myxococcus* cells and *E. coli* cells in the prey colony across time (Figure 5c, see methods). In this assay, we observed that WT *Myxococcus* cells completely take over the *E. coli* population after 72h (Figure 5c). In contrast, the *E. coli* population remained fully viable when in contact with the *kilACF* triple mutant, even after 72h (Figure 5c). Predatory-null phenotypes were also observed in absence of selected Tad structural components, including the secretin (KilC), the ATPase (KilF) and the IM platform protein (KilH) (Figure 5d). A partial defect was observed in the pilin (KilK) but a triple pilin deletion mutant (*kilKLM*) was however completely deficient (Figure 5d).

**Figure 5.**
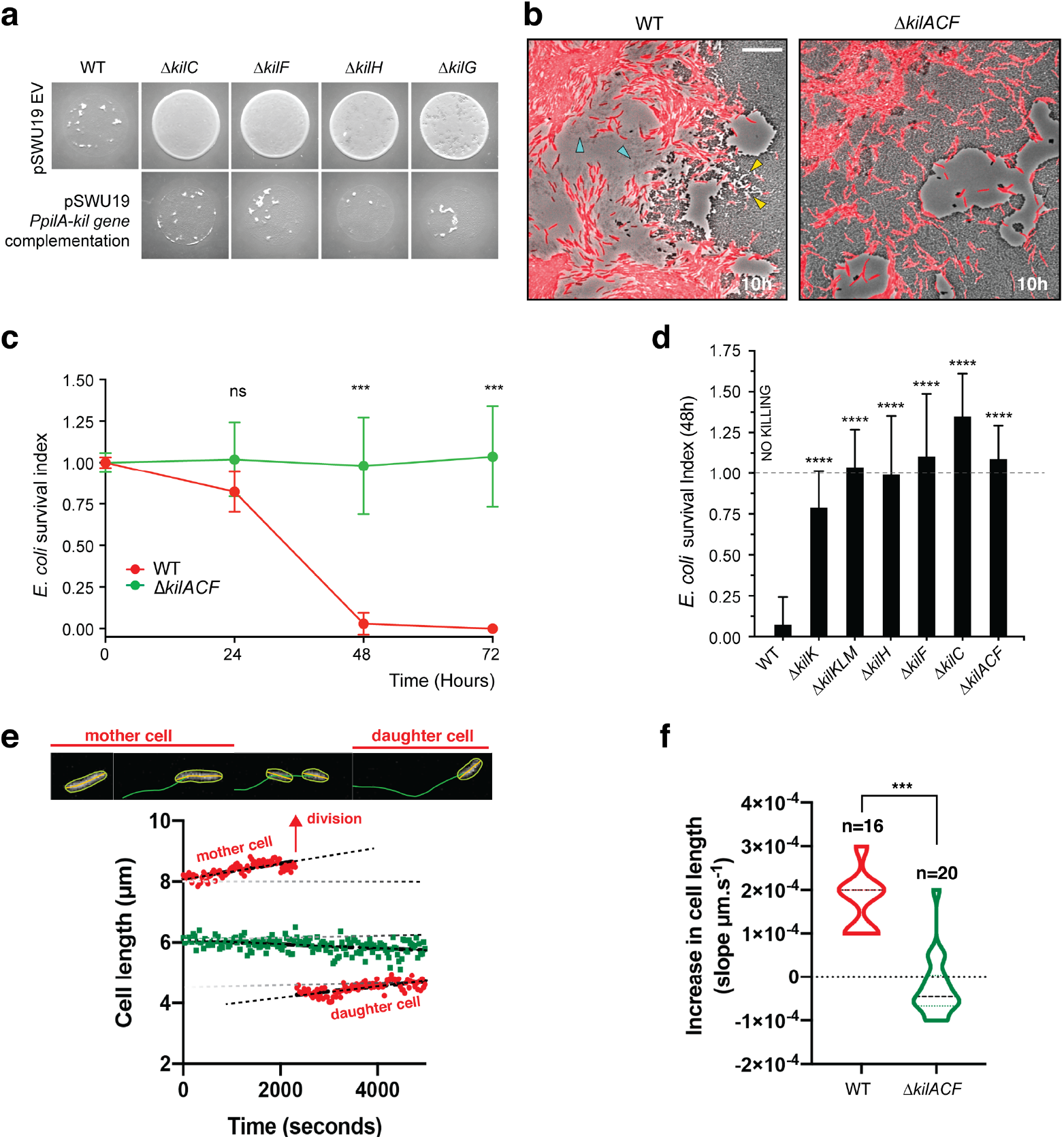
The *kil* genes are required for *M. xanthus* nutrition over prey cells. **a:** The Kil system is essential for predation. Core deletion mutants in Tad-like genes, *kilC* (Secretin), *kilF* (ATPase), *kilH* (IM platform) and *kilG* (IM platform) were mixed with *E. coli* and spotted on CF agar plates (+ 0.07% glucose). After 24 hours of incubation, the mutants carrying the empty vector (EV) pSWU19 were strongly deficient in predation. The same *kil* mutants ectopically expressing the different *kil* genes under the control of the *pilA* promoter presented a restored predation phenotype similar to the WT-EV control. **b:** A *kil* mutant can invade but cannot lyse *E. coli* prey colonies. mCherry-labeled WT and triple *kilACF* mutant are shown for comparison. Note that invading WT cells form corridors (yellow arrowheads) in the prey colony and ghost *E. coli* cells as well as cell debris (blue arrowheads) are left behind the infiltrating *Myxococcus* cells. In contrast, while the Δ*kilACF* penetrates the prey colony, corridors and prey ghost cells are not observed. Scale bar = 10 µm. See corresponding Movie S9 for the full time lapse. **c:** The *kil* genes are essential for prey killing. *E. coli* mCherry cells were measured by FACS at time 0, 24, 48 and 72 hours after the onset of predation. The *E. coli* survival index was calculated by dividing the percentage of “*E. coli* events” at t= 24, 48 or 72 hours by the percentage of “*E. coli* events” at the beginning of the experiment (t=0). This experiment was performed over two biological replicates, in total 6 samples per time point were collected. For each sample, 500,000 events were analyzed. Each data point indicates the mean ± the standard deviation. For each time point, unpaired t-test (with Welch’s correction) statistical analysis was performed to evaluate if the differences observed, relative to wild-type, were significant (***: p≤0.001) or not (ns: p>0.05). **d:** *E. coli* survival in the various *kil* mutant strains at 48 h. *E. coli* mCherry cells were measured (counted) by FACS at time 0 and 48 h after predation. This experiment was performed over three biological replicates, n= 9 per strain and time point. Events were counted as a) and each data point indicates the mean ± the standard deviation. One-way ANOVA statistical analysis followed by Dunnett’s posttest was performed to evaluate if the differences observed, relative to wild-type, were significant (****: p≤0.0001). **e, f:** The *kil* genes are essential for *Myxococcus* growth on prey. **e:** cell growth during invasion. Cell length is a function of cell age during invasion and can be monitored over time in WT cells (in red). In contrast, cell length tends to decrease in a Δ*kilACF* mutant (in green) showing that they are not growing in presence of prey. See associated Movie S10 for the full time lapse. **f:** Quantification of cell growth in WT and Δ*kilACF* mutant backgrounds. Each individual cell was tracked for 5 hours in two biological replicates for each strain. Violin plot of the growth distributions (shown as the cell size increase slopes) are shown. Statistics: Student t-test, ***: p<0.001.

To further test whether Kil-dependent prey killing provides the necessary nutrient source, we directly imaged *Myxococcus* growth in prey colonies, tracking single cells over the course of 6 hours (see methods). This analysis revealed that invading *Myxococcus* cell grow actively during prey invasion. The *Myxococcus* cell cycle could be imaged directly in single cells: cell size increased linearly up to a certain length, which was followed by a motility pause and cytokinesis (Figure 5e, Movie S10). The daughter cells immediately resumed growth at the same speed (Figure 5e). Cell size and cell age are therefore linearly correlated allowing estimation of a ∼5.5 hours generation time from a compilation of traces (Figure 5f, n=16). When the Δ*kilACF* mutant was similarly observed, cell size tended to decrease with time and cell division was not observed (Figures 5e-5f, n=20). Cell shortening could be a consequence of starvation, as observed for example in *Bacillus subtilis*^27^ (although this remains to be documented in *Myxococcus*). Taken together, these results demonstrate the central function of the Kil Tad apparatus in prey killing and consumption.

### The Kil system promotes killing of phylogenetically diverse prey bacteria

*Myxococcus* is a versatile predator and can attack and digest a large number of preys^28,29^. We therefore tested if the Kil system also mediates predation by contact-dependent killing of other bacterial species. To this aim, we tested evolutionarily-distant preys, diderm bacteria, *Caulobacter crescentus, Salmonella typhimurium* and *Pseudomonas aeruginosa*, and monoderm, *Bacillus subtilis*. In plate assays, *M. xanthus* was able to lyse all tested preys, except *P. aeruginosa* (Figure 6a-f). When the Kil system was deleted, the predation ability of *M. xanthus* was severely diminished in all cases (Figure 6a-f). Consistently, *Myxococcus* assembled NG-KilD clusters in contact with *Caulobacter, Salmonella* and *Bacillus* cells, which in all cases led to cell plasmolysis (Figure 6g-i, Movie S11-13). *Myxococcus* cells were however unable to form lethal clusters when mixed with *Pseudomonas* cells (Movie S14), suggesting that, although the Kil system has a large spectrum of target species, it is not universally effective and resistance/evasion mechanisms must exist.

**Figure 6:**
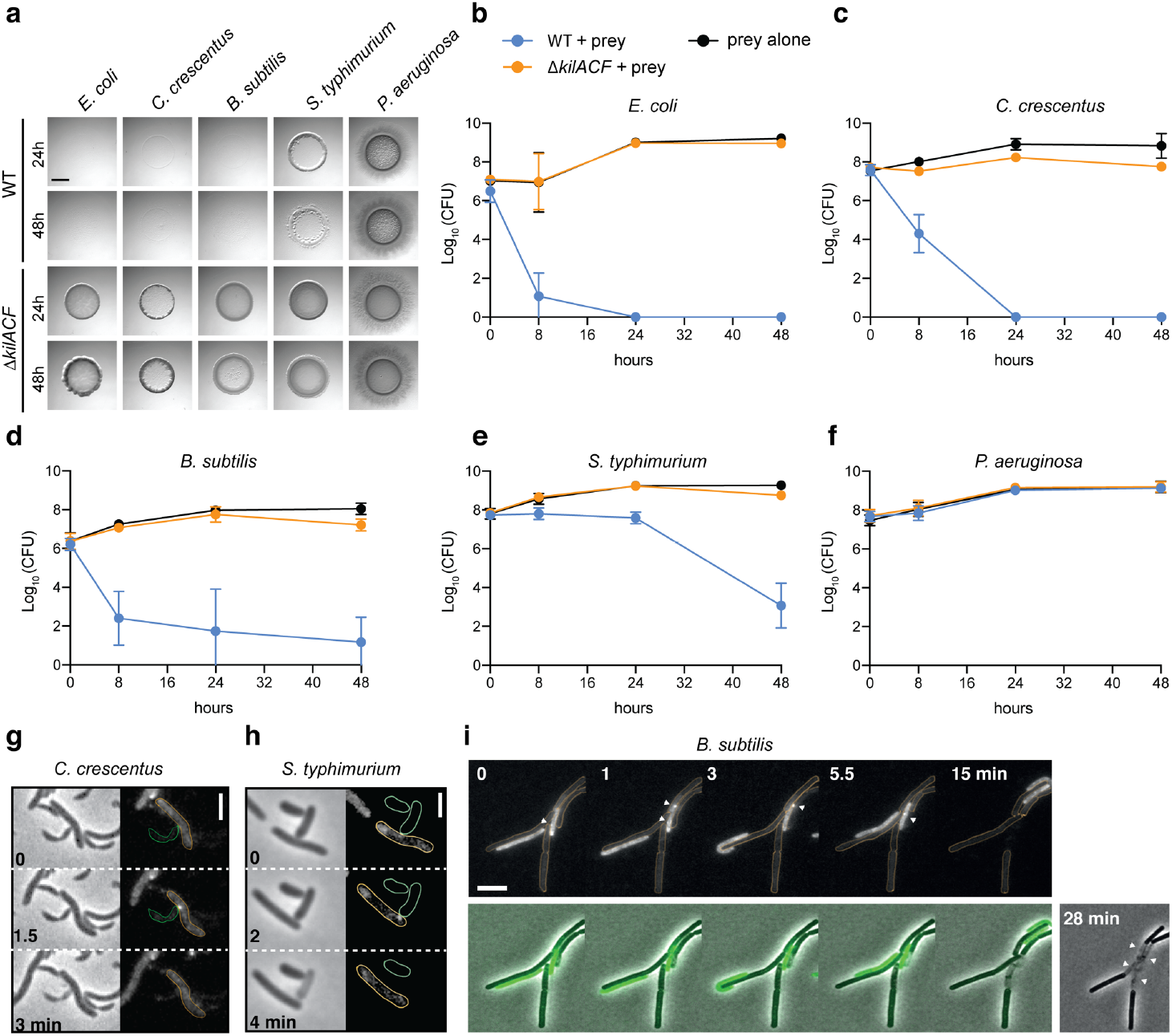
The Kil system mediates killing against diverse bacterial species. **a:** The *kil* genes are predation determinants against various species. To evaluate if *M. xanthus kil* mutant had lost the ability to lyse by direct contact different preys, prey-cell suspensions were directly mixed with *M. xanthus* WT or Δ*kilACF* and spotted on CF agar (+ 0.07% glucose). After 24 and 48 hours of incubation, pictures of the spots corresponding to the different predator/prey couples were taken. Note that *Pseudomonas aeruginosa* is resistant in this assay. **b, c, d, e, f:** Prey cell survival upon predation was evaluated by CFU counting. The different preys were mixed with *M. xanthus* WT (blue circles) or Δ*kilACF* (orange circles) strains and spotted on CF agar (+ 0.07% glucose). Spots were harvested after 0, 8, 24 and 48 hours of predation, serially diluted and platted on agar plates with kanamycin for CFU counting. The prey alone (black circles) was used as a control. Two experimental replicates were used per time point. This experiment was independently performed three times. Error bars represent the standard deviation to the mean. **g:** NG-KilD cluster formation and subsequent contact-dependent killing of *Caulobacter crescentus*. Scale bar = 2 µm. See corresponding Movie S11 for the full time lapse. **h:** NG-KilD cluster formation and subsequent contact-dependent killing of *Salmonella enterica* Typhimurium. Scale bar = 2 µm. See corresponding Movie S12 for the full time lapse. **i:** NG-KilD cluster formation in contact with *B. subtilis*. See corresponding Movie S13 for the full time lapse. Scale bar = 2 µm.

### The kil genes evolved in predatory bacteria

We next explored bacterial genomes for the presence of *kil*-like genes. Phylogenetic analysis indicates that the ATPase (KilF), IM platform proteins (KilH and KilG) and CpaB protein (KilB) share similar evolutionary trajectories, allowing the construction of a well-supported phylogenetic tree based on a supermatrix (Figure 7, see methods). This analysis reveals that Kil-like systems are indeed related to Tad systems (ie Tad systems from alpha-proteobacteria, Figure 7) but they form specific clades in deltaproteobacteria, specifically in *Myxococcales*, in *Bdellovibrionales* and in the recently discovered *Bradymonadales*. In these bacteria, predicted Kil machineries are very similar to the *Myxococcus* Kil system, suggesting a similar function (Figure 7, Table S2). Remarkably, these bacteria are all predatory; the predatory cycle of *Bradymonadales* is yet poorly described but it is thought to be quite similar to the *Myxococcus* predatory cycle, involving surface motility and extracellular prey attack^1^. At first glance, *Bdellovibrio* species use a distinct predatory process, penetrating the prey cell to actively replicate in their periplasmic space^2^. However, this cycle involves a number of processes that are similar to Myxobacteria: *Bdellovibrio* cells also attack prey cells using gliding motility^30^ and attach to them using Type IV pili and a number of common regulatory proteins^31^. Prey cell penetration follows from the ability of the predatory cell to drill a hole into the prey PG at the attachment site^32^. While there is currently no direct evidence that the *Bdellovibrio* Kil-like system is involved in this process, multiple genetic evidence suggest that the Kil homolog are important for prey invasion and attachment^33,34^. It is therefore possible that acquisition of a Tad-like system in deltaproteobacteria was key to the emergence of predation, following its specialization in a possible ancestor of the *Myxococcales, Bdellovibrionales* and *Bradymonadales*.

**Figure 7.**
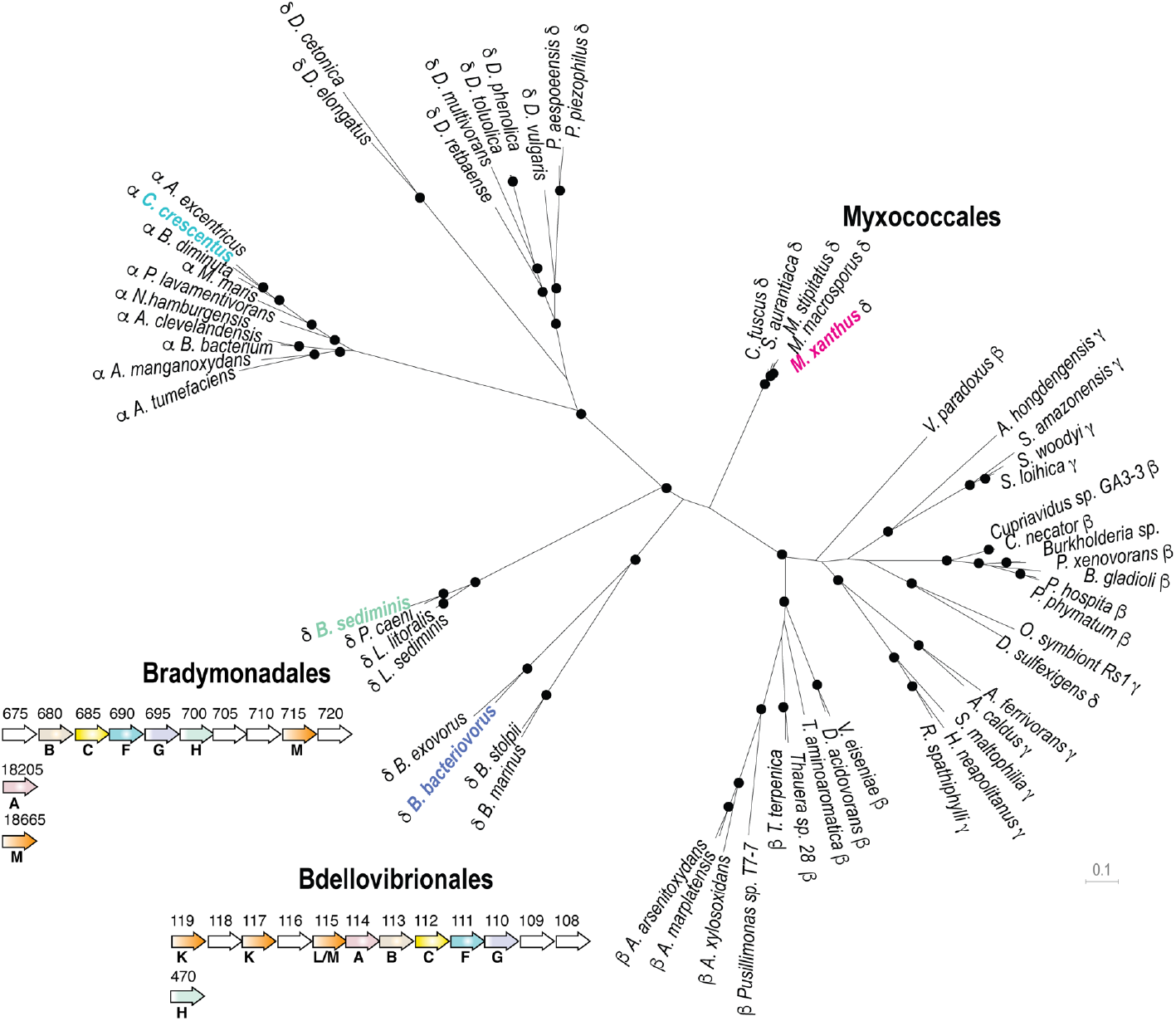
The Kil system is conserved in predatory delta-proteobacteria. Phylogenetic tree of the Type-IV filamentous system that gave rise to the *M. xanthus* Kil system. Only the 4 well-conserved Kil system components were used for constructing the phylogenetic tree. Dots indicate stable bootstrap values (> 75), classes are indicated next to species names. The *M. xanthus* Kil system is also found in other *Myxococcales* and closely related systems are also present in *Bradymonadales* and *Bdellovibrionales*, suggesting a functional specialization related to predation. The genetic organization of *kil*-like genes is shown for example members of each orders, *Bradymonas sediminis* and *Bdellovibrio bacteriovorus* (see also Table S2). The nomenclature and color code for Kil homologs are the same as in Figure 3. Gene accession numbers (KEGG) are shown above gene symbols.

## Discussion

Prior to this work, *Myxococcus* predation was thought to be multifactorial and involve motility, secreted proteins, Outer Membrane Vesicles (OMVs) and antibiotics (ie Myxovirescin and Myxoprincomide) to kill and digest preys extracellularly^3,5^. While a contribution of these processes is not to be ruled out, most likely for prey cell digestion rather than killing (for example by degradative enzymes^3^), we show here that contact-dependent killing is the major prey killing mechanism. In *Myxococcus*, contact-dependent killing can be mediated by several processes, now including T6SS, OME and Kil. We exclude a function for the T6SS, for which a function in *Myxococcus* interspecies interactions has yet to be demonstrated. Rather, it appears that together with OME, Type VI secretion controls a phenomenon called social compatibility, in which the exchange of toxins between *Myxococcus* cells prevents immune cells from mixing with non-immune cells^19^. We have not tested a possible function of OME in prey killing because OME allows transfer of OM protein and lipids between *Myxococcus* cells when contact is established between identical outer membrane receptors, TraA^9^. OME is therefore highly *Myxococcus* species and even strain-specific and mediates social compatibility when SitA lipoprotein toxins are delivered to non-immune TraA-carrying *Myxococcus* target cells^35^.

The Kil system is both required for contact-dependent killing in liquid and on surfaces. Remarkably, proteins belonging to each motility systems show distinct requirement in liquid or on solid media. In liquid, Type-IV pili mediate prey-induced biofilm formation, which likely brings *Myxococcus* in close contact with the prey cells. This intriguing process likely requires EPS (since *pilA* mutants also lack EPS^36^), which deserves further exploration. On surfaces, likely a more biologically relevant context, contact-dependent killing is coupled to A-motility to penetrate prey colonies and interact with individual prey cells. The prey recognition mechanism is especially intriguing because dynamic assembly of a Tad-like system at the prey contact site is a novel observation; in general, these machineries and other Type IV filamentous systems^22^, such as TFPs tend to assemble at fixed cellular sites, often a cell pole^11,23,37^. KilD clusters do not require a functional Tad-like system to form in contact with prey cells, suggesting prey contact induces Tad assembly via an upstream signaling cascade. Such sensory system could be encoded within the clusters 1 and 2, which contain a large number of conserved genes with unknown predicted functions (up to 11 proteins of unknown functions just considering cluster 1 and 2, Figure 3a, Table S1). In particular, the large number of predicted proteins with FHA^24^ type domains (Table S1) suggests a function in a potential signaling cascade. In *Pseudomonas aeruginosa*, FHA domain-proteins act downstream from a phosphorylation cascade triggered by contact, allowing *Pseudomonas* to fire its T6SS upon contact^38^. This mechanism is triggered by general perturbation of the *Pseudomonas* membrane^39^, which could also be the case for the Kil system. Kil assembly is provoked both by monoderm and diderm bacteria, which suggests that prey-specific determinants are unlikely. Recognition is nevertheless non-universal and does not occur in contact with *Pseudomonas* or *Myxococcus* itself. Therefore, evasion mechanisms must exist, perhaps in the form of genetic determinants that shield cells from recognition.

The Kil Tad-like system itself is required to pause A-motility and prey cell killing. Motility regulation could be indirect because differential effects are observed depending on *kil* gene deletions (Figures 3 and 4), suggesting that assembly of a functional Tad apparatus is not strictly required for regulation. In contrast, prey killing requires a functional Tad apparatus. In particular, the pilin proteins are required during prey invasion but they are dispensable (partially) in liquid cultures showing that they do not promote toxicity. In liquid, direct contacts may be enforced by TFPs in the biofilm, perhaps rendering the Tad pilins partially redundant. Tad Pilin function would however become essential to onto prey cells on surfaces where the pili are dispensable. How the pilins organize to form polymers and whether they do, remains to be determined; the lack of the major pilin (KilK) is compensated by the remaining pseudo-pilins KilL and M, which is somewhat surprising given that pseudopilins are generally considered to prime assembly of major pilin polymers^22^. It is currently unclear if the Kil system is also a toxin-secretion device; for example, if it also functioned as a Type II secretion system. Alternatively, the Kil complex might recruit a toxin delivery system at the prey contact site. This latter hypothesis is in fact suggested by the remaining low (but still detectable) contact-dependent toxicity of *kil* mutants (Figures 3 and 4). Given that *Myxococcus* induces prey PG degradation locally, we hypothesize that a secreted cell wall hydrolase becomes active at the prey contact site. This is not unprecedented: *Bdellovibrio* cells secrete a sophisticated set of PG modifying enzymes, D,D-endopeptidases^40^, L,D transpeptidases^32^ and Lysozyme-like enzymes^41^ to penetrate prey cells, carve them into bdelloplasts and escape. In *Myxococcus*, deleting potential D,D-endopeptidases^42^ (Δ*dacB*) did not affect predation (Figure S5) which might not be surprising given that *Myxococcus* simply lyses its preys while *Bdellovibrio* needs to penetrate them while avoiding their lysis to support its intracellular cycle. The *Myxococcus* toxin remains to be discovered, bearing in mind, that similar to synergistic toxic T6SS effectors^43^, several toxic effectors could be injected, perhaps explaining how *Myxococcus* is able to kill both monoderm and diderm preys.

The Myxobacteria are potential keystone taxa in the soil microbial food web^44^, meaning that Kil-dependent mechanisms could have a major impact in shaping soil eco-systems. While the Kil proteins are most similar to proteins from Tad systems, there are a number of key differences that suggest profound diversification: (i), the Kil system involves a single ATPase and other Tad proteins such as assembly proteins TadG, RcpB and pilotin TadD are missing^22^; (ii), several Kil proteins have unique signatures, the large number of associated genes of unknown function; in particular, the over-representation of associated FHA domain proteins, including the central hexameric ATPase KilF itself fused to an N-terminal FHA domain. The KilC secretin is also uniquely short and lacks the N0 domain, canonically found in secretin proteins^45^, which could be linked to increased propensity for dynamic recruitment at prey contact sites. Future studies of the Kil machinery could therefore reveal how the contact-dependent properties of Tad pili were adapted to prey cell interaction and intoxication, likely a key evolutionary process in predatory bacteria.

## Methods

### Bacterial strains, growth conditions, motility plates, western blotting and genetic constructs

See Tables S3-S5 for strains, plasmids, and primers. *E. coli* cells were grown under standard laboratory conditions in Luria-Bertani (LB) broth supplemented with antibiotics, if necessary. *M. xanthus* strains were grown at 32°C in CYE (Casitone Yeast Extract) rich media as previously described^42^. *S. enterica* Typhimurium, *B. subtilis* and *P. aeruginosa* were grown overnight at 37°C in LB. *C. crescentus* strain NA1000 was grown overnight at 30°C in liquid PYE (Peptone Yeast Extract). Motility plate assays were conducted as previously described on soft (0.3%) or hard (1.5%) agar CYE plates^46^.

The deletion strains and the strains expressing the different Neon Green fusions were obtained using a double-recombination strategy as previously described^46 47^. Briefly, the *kil* deletion alleles (carrying ∼700-nucleotide long 5’ and 3’ flanking sequences of *M. xanthus* locus tags) were Gibson assembled into the suicide plasmid pBJ114 (*galK*, Kan^R^) and used for allelic exchange. Plasmids were introduced in *M. xanthus* by electroporation. After selection, clones containing the deletion alleles were identified by PCR. Using the same strategy, “Neon Green fusion” alleles were introduced at *kilD* and *kilF* loci. The corresponding strains expressed, under the control of their native promoters, C-terminal Neon Green fusions of KilD and KilF.

For complementation of Δ*kilC*, Δ*kilF*, Δ*kilG* and Δ*kilH* strains, we used the pSWU19 plasmid (Kan^R^) allowing ectopic expression of the corresponding genes from the *pilA* promoter at Mx8-att site. The same strains transformed with the empty vector were used as controls.

To express KilG C-terminally fused to Neon Green, a pSWU19-*PpilA-kilG-NG* was created and transformed in the Δ*kilG* strain.

Western blotting was performed as previously described^46^ using a commercial polyclonal anti Neon-Green antibody (Chromotek).

### Growth in liquid cultures

To compare growth rates of *M. xanthus* WT and Δ*kilACF* strains, overnight CYE cultures were used to inoculate 25 ml of CYE at OD_600_= 0.05. Cultures were then incubated at 32°C with a shaking speed of 160 rpm. To avoid measuring cell densities at night, a second set of cultures were inoculated 12 hours later at OD_600_= 0.05. Every 4 hours, 1 ml sample of each culture was used to measure optical densities at 600 nm with a spectrophotometer. The different measurements were then combined into a single growth curve. This experiment was performed with three independent cultures per strain.

### Predation assay on agar plates

#### Prey colony invasion on CF agar plates

*M. xanthus* and the different prey cells (except *C. crescentus*) were respectively grown overnight in 20 ml of CYE at 32°C and in 20 ml of LB at 37°C. *C. crescentus* was grown in 20 ml of PYE at 30°C. The next day, cells were pelleted and resuspended in CF medium (MOPS 10 mM pH 7.6; KH_2_PO_4_ 1 mM; MgSO4 8 mM; (NH_4_)_2_SO_4_ 0.02%; Na citrate 0.2%; Bacto Casitone 0.015%) to a final OD_600_ of 5. 10 µl of *M. xanthus* and prey cell suspensions were then spotted next to each other (leaving less than 1-mm gap between each spot) on CF 1.5% agar plates with or without 0.07% glucose (to allow minimal growth of the prey cells) and incubated at 32°C. After 48-hours incubation, pictures of the plates were taken using a Nikon Olympus SZ61 binocular loupe (10x magnification) equipped with a camera and an oblique filter. ImageJ software was used to measure the surface of the prey spot lysed by *M. xanthus*.

#### Spotting predator-prey mixes on CF agar plates

to force the contact between *M. xanthus* and a prey, mixes of predator/prey were made and spotted of CF agar plates. 200 µl of a prey cell suspension (in CF, OD_600_ = 5) were mixed with 25 µl of a *M. xanthus* cell suspension (in CF, OD_600_ = 5) and 10 µl of this mix were spotted on CF agar plates supplemented with 0.07% glucose. As described above, pictures of the plates were taken after 24-hour incubation.

### Microscope invasion predation assay and contact-dependent killing

#### Prey colony invasion on CF agar pads

prey invasion was imaged by microscopy using the Bacto-Hubble system (the specific details of the Method are described elsewhere^10^). Briefly, cell suspensions concentrated to OD_600_=5 were spotted at 1 mm distance onto CF 1.5% agar pads and a Gene Frame (Thermo Fisher Scientific) was used to sandwich the pad between the slide and the coverslip and limit evaporation of the sample. Slides were incubated at 32°C for 6 hours before imaging, allowing *Myxococcus* and *E. coli* to form microcolonies. Time-lapse of the predation process was taken at 40x or 100x magnification. Movies were taken at the invasion front where *Myxococcus* cells enter the *E. coli* colony. To facilitate tracking, *M. xanthus* cells were labeled with fluorescence^48^. Fluorescence images were acquired every 30 seconds for up to 10 hours, at room temperature.

#### Spotting predator-prey mixes on CF agar pads

to image contact-dependent killing between *M. xanthus* and prey cells (*E. coli, C. crescentus, B. subtilis, S. typhimurium and P. aeruginosa*), cells were grown as described above, pelleted and resuspended in CF medium to a final OD_600_ of 1. Equal volumes of *M. xanthus* and prey cell suspensions were then mixed together and 1 µl of the mix was spotted on a freshly made CF 1.5% agar pad on a microscope slide. After the spot has dried, the agar pad was covered with a glass coverslip, and incubated in the dark at room temperature for 20-30 minutes before imaging.

Time-lapses experiments were performed using two automated and inverted epifluorescence microscope: a TE2000-E-PFS (Nikon), with a ×100/1.4 DLL objective and an ORCA Flash 4.0LT camera (Hamamatsu) or a Ti Nikon microscope equipped with an ORCA Flash 4.0LT camera (Hamamatsu). Theses microscopes are equipped with the “Perfect Focus System” (PFS) that automatically maintains focus so that the point of interest within a specimen is always kept in sharp focus at all times, despite any mechanical or thermal perturbations. Images were recorded with NIS software from Nikon. All fluorescence images were acquired with appropriate filters with a minimal exposure time to minimize photo-bleaching and phototoxicity effects: 30-minute long time-lapses (one image acquired every 30 seconds) of the predation process were taken at 100x magnification. DIA images were acquired using a 5 ms light exposure and GFP fluorescent images were acquired using a 100 ms fluorescence exposure with power intensity set to 50% (excitation wavelength 470 nm) to avoid phototoxicity.

### Labelling *E. coli* cells with the fluorescent D-Amino Acid TADA

Lyophilized TADA (MW = 381.2g/mol, laboratory stock^15^) was re-suspended in DMSO at 150 mM and conserved at −20°C. The labeling was performed, for 2 h in the dark at room temperature, using 2 μl of the TADA solution for 1ml of cells culture (OD_600_ = 2). Cells were then washed four times with 1ml of CF and directly used for predation assays on agar pad.

### Image Analysis

Image analysis was performed under FIJI^49^ and MicrobeJ^50^ an ImageJ plug-in for the analysis of bacterial cells.

#### Semantic segmentation of *Myxococcus* cells

was obtained using the newly developed MiSiC system, a deep learning based bacterial cell segmentation tool^10^. The system was used in semantic segmentation mode and annotated manually to reveal *E. coli* lysing cells.

#### Kymograph construction

Kymographs were obtained after manual measurements of fluorescence intensities along FIJI hand-drawn segments and the FIJI-Plot profile tool. The measurements were then exported into the Prism software (Graphpad, Prism 8) to construct kymographs.

#### Cell tracking

Cell tracking and associated morphometrics were obtained using MicrobeJ. Image stacks were first processed stabilized and filtered with a moderate Gaussian blur and cells were detected by thresholding and fitted with the Plug-in “medial axis” model. Trajectories were systematically verified and corrected by hand when necessary.

#### Tracking *Myxococcus* pauses in contact with a prey, NG-KilD foci formation and prey cell lysis

in 30-minute time-lapses, contacts between prey cells and *Myxococcus* cells were scored. Pauses were counted when the predatory cell stopped all movement upon contact with the prey. We also counted if these contacts lead to the formation of NG-KilD foci and to cell lysis. Thus, for a determined *E. coli* cell, we scored the number of contacts with *Myxococcus*, the number of pauses these contacts induces in *M. xanthus* motility, the number of NG-KilD foci formed upon contacts and, ultimately, the lysis of the cell. Five independent movies were analyzed for each strain and the percentage of contacts leading to a pause in motility, NG-KilD foci formation and cell lysis was calculated. We also estimated the percentage of NG-KilD clusters leading to cell lysis

#### Tracking cluster time to lysis

Time to lysis measures the elapsed time between cluster appearance to prey cell death. Data were obtained from two biological replicates.

### CPRG assay for contact-dependent killing in liquid

#### CPRG assay in 24-well plates

*M. xanthus* and *E. coli* cultures were grown overnight, pelleted and resuspended in CF at OD_600_ ∼5. 100 µl of *M. xanthus* cell suspension (WT and mutants) were mixed with 100 µl of *E. coli* cell suspension in a 24-well plate containing, in each well, 2 ml of CF medium supplemented with CPRG (Sigma Aldrich, 20 µg/ml) and IPTG (Euromedex, 50 µM) to induce *lacZ* expression. The plates were then incubated at 32°C with shaking and pictures were taken after 24 and 48 hours of incubation. To test the contact-dependance, a two-chamber assay was carried out in a Corning 24 well-plates containing a 0.4-µm pore polycarbonate membrane insert (Corning Transwell 3413). This membrane is permeable to small metabolites and proteins and impermeable to cells. *E. coli* cells were inoculated into the top chamber and *M. xanthus* cells into the bottom chamber.

#### CPRG assay in 96-well plates

To evaluate the predation efficiency of the different *kil* mutants, the CPRG assay was adapted as follow: wild-type *M. xanthus* and the *kil* mutant strains were grown overnight in 15 ml of CYE. *E. coli* was grown overnight in 15 ml of LB. The next morning, *M. xanthus* and *E. coli* cells were pelleted and resuspended in CF at OD_600_ = 0.5 and 10, respectively. To induce expression of the β-galactosidase, IPTG (100 µM final) was added to the *E. coli* cell suspension.

In a 96-well plate, 100 µl of *M. xanthus* cell suspension were mixed with 100 µl of *E. coli* cell suspension. Wells containing only *M. xanthus, E. coli* or CF were used as controls. The lid of the 96-well plate was then sealed with a breathable tape (Greiner bio-one) and the plate was incubated for 24 hours at 32°C while shaking at 160 rpm. In this setup, we observed that *M. xanthus* and *E*.*coli* cells aggregate at the bottom of the well and therefore come in direct contact, favoring predation in liquid.

The next day, the plate was centrifuged 10 minutes at 4800 rpm and 25 µl of the supernatant were transferred in a new 96-well plate containing 125 µl of Z-buffer (Na_2_HPO_4_ 60 mM, NaH_2_PO_4_ 40 mM, KCl 10 mM pH7) supplemented with 20 µg/ml of CPRG. After 15-30 minutes of incubation at 37°C, the enzymatic reaction was stopped with 65 µl of Na_2_CO_3_ (1 M) and the absorbance at 576 nm was measured using a TECAN Spark plate reader.

This experiment was performed independently four times. For Miller unit calculation, after absorbance of the blank (with CF) reaction was subtracted, the absorbances measured at 576 nm were divided by the incubation time and the volume of cell lysate used for reaction. The resulting number was then multiplied by 1000.

### Crystal violet biofilm staining

In a 96-well plate, 100 µl of *M. xanthus* cell suspension (in CF, OD_600_ = 0.5) were mixed with 100 µl of *E. coli* cell suspension (in CF, OD_600_ = 10) and incubated for 24 hours at 32°C while shaking at 160 rpm. The next day, the supernatant was carefully removed and the wells were washed with 200 µl of CF twice. Then, 100 µl of a 0.01% crystal violet solution were added to each well and incubated for 5 minutes. Wells were washed twice with 200 µl of water before imaging.

### Prey CFU counting after predation

*E. coli, S. typhi, P. aeruginosa* and *B. subtilis* kanamycin resistant strains were grown at 37°C in liquid LB supplemented with kanamycin (50 or 10 µg/ml). *C. crescentus* kanamycin resistant strain was grown at 30°C in liquid PYE supplemented with kanamycin (25 µg/ml). Wild-type and Δ*kilACF M. xanthus* strains were grown at 32°C in liquid CYE. Cells were then centrifuged and pellets were resuspended in CF at an OD_600_ of 5. 25 µl of *M. xanthus* cell suspension and 200 µl of prey cell suspension were then mixed together and 10 µl were spotted on CF agar plates supplemented with 0.07% glucose. After drying, plates were incubated at 32°C. At 0, 8, 24, and 48-hour time points, spots were harvested with a loop and resuspended in 500 µl of CF. This solution was then used to make 10-fold serial dilutions in a 96-well plate containing CF. At the exception of *C. crescentus*, 5 µl of each dilution were spotted on LB agar plates supplemented with 10 µg/ml of kanamycin and incubated at 37°C for 24 hours. *C. crescentus* dilutions were spotted on PYE agar plates supplemented with 25 µg/ml of kanamycin and incubated at 30°C for 24 hours. The next day, colony-forming units were counted and the number of prey cells that survived in the predator/prey spot was calculated.

### Fluorescence-Activated Cell Sorting (FACS) measurements of *E. coli* killing

*M. xanthus* strains (wild-type and *kil* mutants) constitutively expressing GFP were grown overnight in liquid CYE without antibiotics. *E. coli* mCherry (prey) was grown overnight in liquid LB supplemented with ampicillin (100 µg/ml). The next morning, optical densities of the cultures were adjusted in CF medium to OD_600_= 5. *M. xanthus* GFP and *E. coli* mCherry cell suspensions were then spotted onto fresh CF 1.5% agar plates as previously described^46^. Briefly, 10-µl drops of the prey and the predator cell suspensions were placed next to each other and let dry. Inoculated plates were then incubated at 32°C. Time 0 corresponds to the time at which the prey and the predator spots were set on the CF agar plate. At time 0, 24, 48 and 72 hours (post predation) and for each *M. xanthus* strain, two predator/prey spot couples were harvested with a loop and resuspended in 750 µl of TPM. To fix the samples, paraformaldehyde (32% in distilled water, Electron Microscopy Sciences) was then added to the samples to a final concentration of 4%. After 10-min incubation at room temperature, samples were centrifuged (8 min, 7500 rpm), cell pellets were then resuspended in fresh TPM and optical densities were adjusted to OD_600_ ∼0.1.

Samples were then analyzed by flow cytometry. Flow cytometry data were acquired on a Bio-Rad S3e Cell Sorter and analyzed using the ProSort software, version 1.6. For each sample, a total population of 500,000 events was used and events corresponding to the sum of *M. xanthus*-GFP and *E. coli*-mCherry. A blue laser (488 nm, 100mW) was used for detection of forward scatter (FSC) and side scatter (SSC) and for excitation of GFP. A yellow-green laser (561 nm, 100 mW) was used for excitation of mCherry. GFP and mCherry signals were collected using, respectively, the emission filters FL1 (525/30 nm) and FL3 (615/25 nm) and a compensation was applied on the mCherry signal. Samples were run using the low-pressure mode (10,000 particles/s). To calibrate the instrument and reduce background noise, suspensions of fluorescent and non-fluorescent *M. xanthus* and *E. coli* cells were used: a threshold was applied on the FSC signal, and voltages of the photomultipliers for FSC, SSC, FL1 and FL3 were also adjusted. The density plots obtained (small angle scattering FSC versus wide angle scattering SSC signal) were first gated on the overlapped population of *M. xanthus* and E. coli, filtered to remove the multiple events and finally gated for high FL1 signal (*M. xanthus*-GFP) and high FL3 signal (*E. coli*-mCherry).

### Bioinformatic analyses

#### Homology search strategy

we used several search strategies to identify all potential homologous proteins of the Kil system: we first used BLAST^51,52^ to search for reciprocal best hits (RBH) between the *M. xanthus* and the *B. bacteriovorus* and *B. Sediminis* Kil systems, as well as the *C. crescentus* Tad system, identifying *bona fide* orthologs between the three species. We limited the search space to the respective proteomes of the three species. We then used HHPRED^53^ to search for remotely conserved homologs in *B. bacteriovorus* using the proteins from the two operons identified in *M. xanthus*. Finally, we performed domain comparisons between proteins from the *B. bacteriovorus* and *B. sediminis* Kil operons and *C. crescentus* Tad system to identify proteins with similar domain compositions in *M. xanthus*. Identified orthologs or homologs between the three species, the employed search strategy, as well as resulting e-values are shown in Table S2.

#### Structure predictions

tertiary structural models of secretin and cytoplasmic ATPase were done using Phyre2^54^or SWISS-MODEL^55^, in both cases using default parameters. Quaternary models were generated using SWISS-MODEL. Structural models were displayed using Chimera^56^ and further processed in Illustrator ™.

#### Phylogenetic analyses

we used the four well-conserved Kil system components for phylogenetic analysis. To collect species with secretion systems similar to the Kil system, we first used MultiGeneBLAST^57^ with default parameters. Orthologs of the four proteins from *B. bacteriovorus, B. Sediminis* and *C. crescentus* from closely related species were added manually. We aligned each of the four proteins separately using MAFFT^58^ and created a supermatrix from the four individual alignments. Gblocks^58^ using relaxed parameters was used prior to tree reconstruction to remove badly aligned or extended gap regions. The resulting alignment is shown in Suppl. File 1. Alignments of individual trees were also trimmed using Gblocks. PhyML^59^ was used for tree reconstruction, using the JTT model and 100 bootstrap iterations. Trees were displayed with Dendroscope^60^ and further processed in Illustrator ™.

## Acknowledgements

We thank Lotte Søgaard-Andersen and Anke Treuner-Lange for the gift of the VipA plasmid. We thank Laurent Aussel for *E. coli* plasmids, Anne Galinier lab for the *Bacillus subtilis* strains, Emanuele Biondi lab for the *Caulobacter crescentus* strains and Sophie Bleves for the *Pseudomonas aeruginosa* strain. We thank Dorothée Murat, Romé Voulhoux, Marcelo Nöllmann, Vladimir Pelicic and Friedhelm Pfeiffer for discussions.

Research in TM lab was supported by a 2019 CNRS 80-Prime allowance on bacterial predation and pattern formation. SS and PDB are supported by an MENRT thesis grant from the ministry of research.

## Author contributions

SS, JH and TM conceived the experiments and analyzed the data. SS, JH and DR performed most experiments. GB ran FACS experiments and analyzed data. PDB and BH performed bioinformatic analysis, homology searches, structure predictions and phylogenetic analysis. LM, EC, SS and TM conceived and analyzed T6SS experiments. RM provided data with the A^-^S^-^ motility mutant. TM and JH wrote the paper.

## Extended Figures and Legends

**Figure S1.**
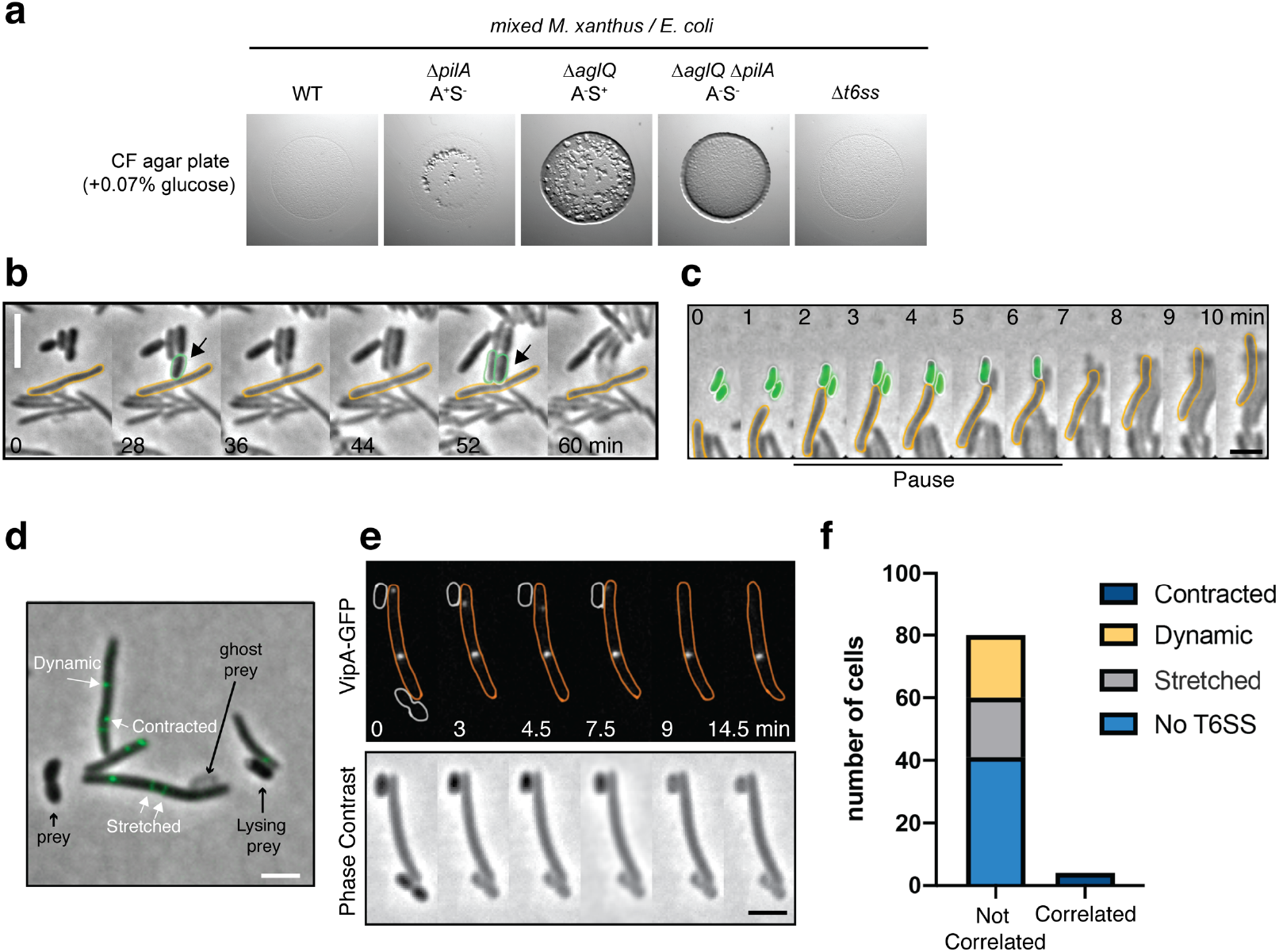
The motility complexes and the Type-6 Secretion System do not mediate contact-dependent killing. **a:** WT *M. xanthus* and the different motility mutant strains were mixed with *E. coli* and spotted on CF 1.5% agar plates (+0.07% glucose). After 24 hours of incubations pictures were taken, showing that WT and Δ*t6ss* had similar predation efficiencies. A A^+^S^-^ (Δ*pilA*) strain presented a predation efficiency slightly reduced. However, the A^-^S^+^ strain (Δ*aglQ*) was greatly impaired at predating. No predation was observed for the A^-^S^-^ strain (Δ*aglQ* Δ*pilA*). Therefore, A-motility appears to be essential for predation. **b:** Contact-dependent killing by an A^-^S^-^ motility mutant (Δ*aglQ* Δ*pilA*). Growth of *E. coli* cells leads to contact with non-motile *Myxococcus* cells and rapid lysis. Example cell reflects events observed for n=20 events. Scale bar = 2 µm. **c:** Contact-dependent killing by a Δ*t6ss* motility mutant. *E. coli* prey cells are labeled with GFP to monitor contact-dependent lysis. Example cell reflects events observed for n=20 events. Scale bar = 2µm. **d-e:** T6SS VipA sheath assembly in *Myxococcus* cells during predation. Several assembly patterns are observed as described in other bacteria. Stretched: extended T6SS sheaths. Contracted: retracted T6SS sheath. Scale bars = 2 µm. **f:** Prey contact-dependent lysis is not correlated to T6SS sheath contraction. Contact-dependent lysis and VipA-GFP dynamics were observed simultaneously. Contraction and lysis at the contacted site were only marginally observed (correlated) suggesting that T6SS intoxication plays a minor role at best in contact-dependent killing.

**Figure S2.**
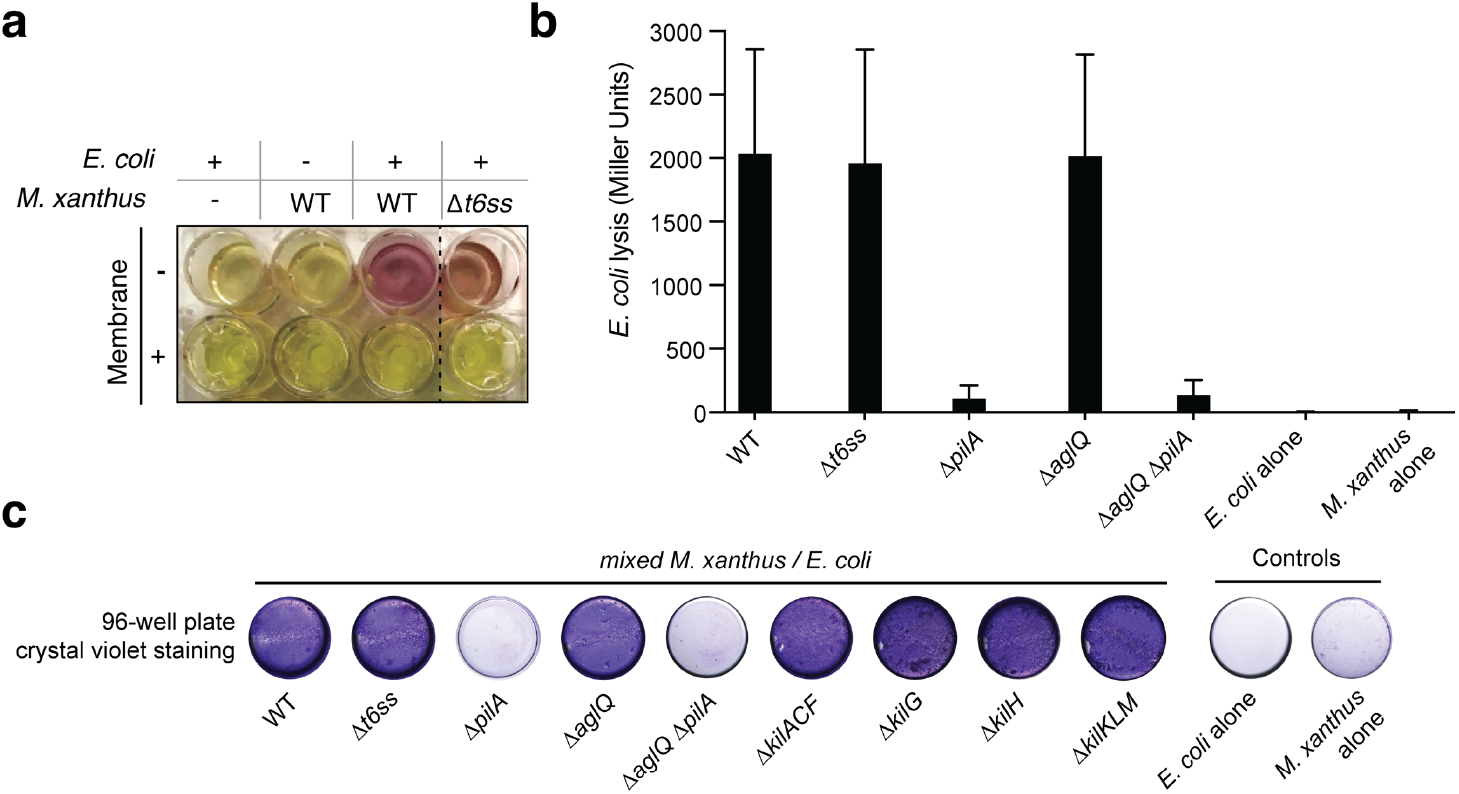
Contact-dependent lysis in liquid cultures. **a:** *E. coli* lysis is detected as extracellular release of LacZ allows hydrolysis of CPRG which becomes colored after 24 hours. Lysis is not observed when *Myxococcus* (WT or Δ*t6ss*) and *E. coli* are separated by a membrane, showing that it is contact-dependent. **b**: A CPRG assay was performed with these same strains as in Figure S1a. β-galactosidase activities of the cell lysates (n=4, expressed in Miller Units) were measured for each strain. After 24-hour incubation, only WT and A^+^S^-^ strains had the ability to lyse *E. coli* in liquid. PilA appears to be essential for cell-cell contact with *E. coli* and cell lysis in liquid. This experiment was independently performed four times. Error bars represent the standard deviation to the mean. **c**: Crystal violet assay. After 24-hour incubation, wells containing the different *M. xanthus* strains mixed with *E. coli* were stained with a crystal violet solution to revealed biofilm formation. The Δ*pilA* strains appeared to be deficient at forming a biofilm in the presence of a prey.

**Figure S3:**
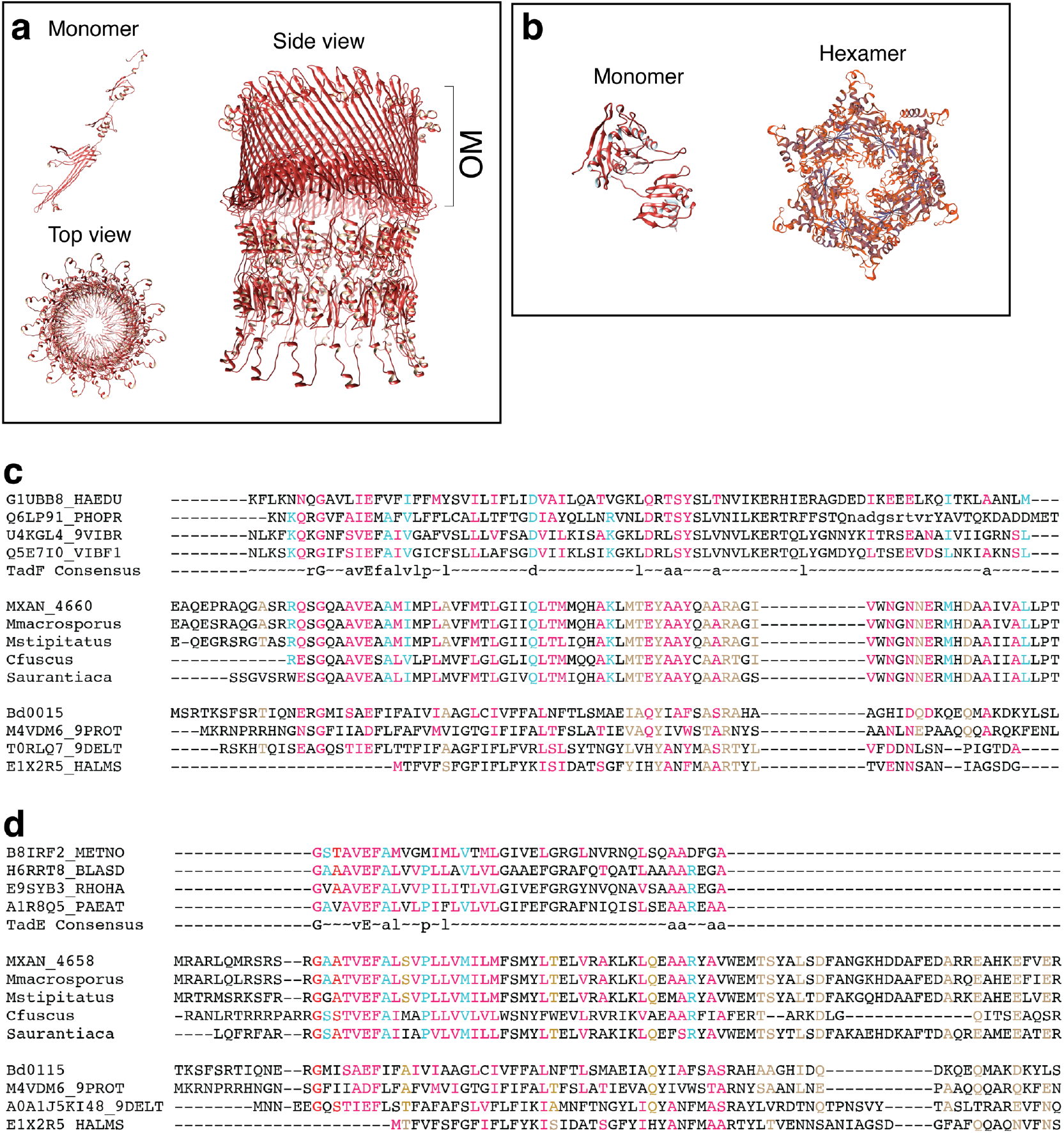
Bioinformatics analyses of Kil proteins. **a, b:** Structural models of the putative KilC secretin (a) and KilF hexameric ATPase (b). KilC Secretin: tertiary and quaternary structural models were based on the structure of *E. coli* type II secretion system GspD protein D (PDB identifier 5WQ7) and generated with SWISS-MODEL (Methods). ATPase: modeled with Phyre2 and SWISS-MODEL based on the structure of the *Sulfolobus acidocaldarius* FlaI ATPase (PDB identifier 4II7). **c**,**d:** Analysis of putative pseudo-pilin proteins. For clarity, the multiple alignment is separated in three blocks representing the three different groups, the alpha-proteobacteria, the *Myxococcales* and the *Bdellovibrionales*. All sequences except the one from *C. fuscus* were taken from HHPRED matrix alignments. Residues conserved between the pfam domains TadF and the MXAN_4660 family, as well as TadE and the MXAN_4658 family, respectively, are highlighted in cyan; those conserved between the Bd0115 family and the MXAN_4660 family, as well as TadE and the MXAN_4658 family, respectively are highlighted in brown; residues conserved in all (TadF, MXAN_4660, Bd0115, TadE, MXAN_4658, Bd0115, respectively) are highlighted in red. **c:** HHPRED-based multiple sequence alignment of MXAN_4660 (KilM) with the TadF domain and pilus assembly protein Bd0115 from *B. bacteriovorus*. Myxobacterial sequences correspond to the following NCBI RefSeq IDs: *Myxococcus xanthus*: WP_011554652; *Myxococcus stipitatus*: WP_015350653; *Myxococcus macrosporus*: WP_043711698; *Stigmatella aurantiaca*: WP_013376800; *Cystobacter fuscus*: WP_002624349. **d:** HHPRED-based multiple sequence alignment of MXAN_4658 with the TadE domain and pilus assembly protein Bd0115 from *B. bacteriovorus*. Myxobacterial sequences correspond to the following NCBI RefSeq IDs: *M. xantus*: WP_011554650; *M. stipitatus*: WP_015350651; *M. macrosporus*: WP_043711696; *S. aurantiaca*: WP_013376798; *C. fuscus*: WP_002624796.

**Figure S4:**
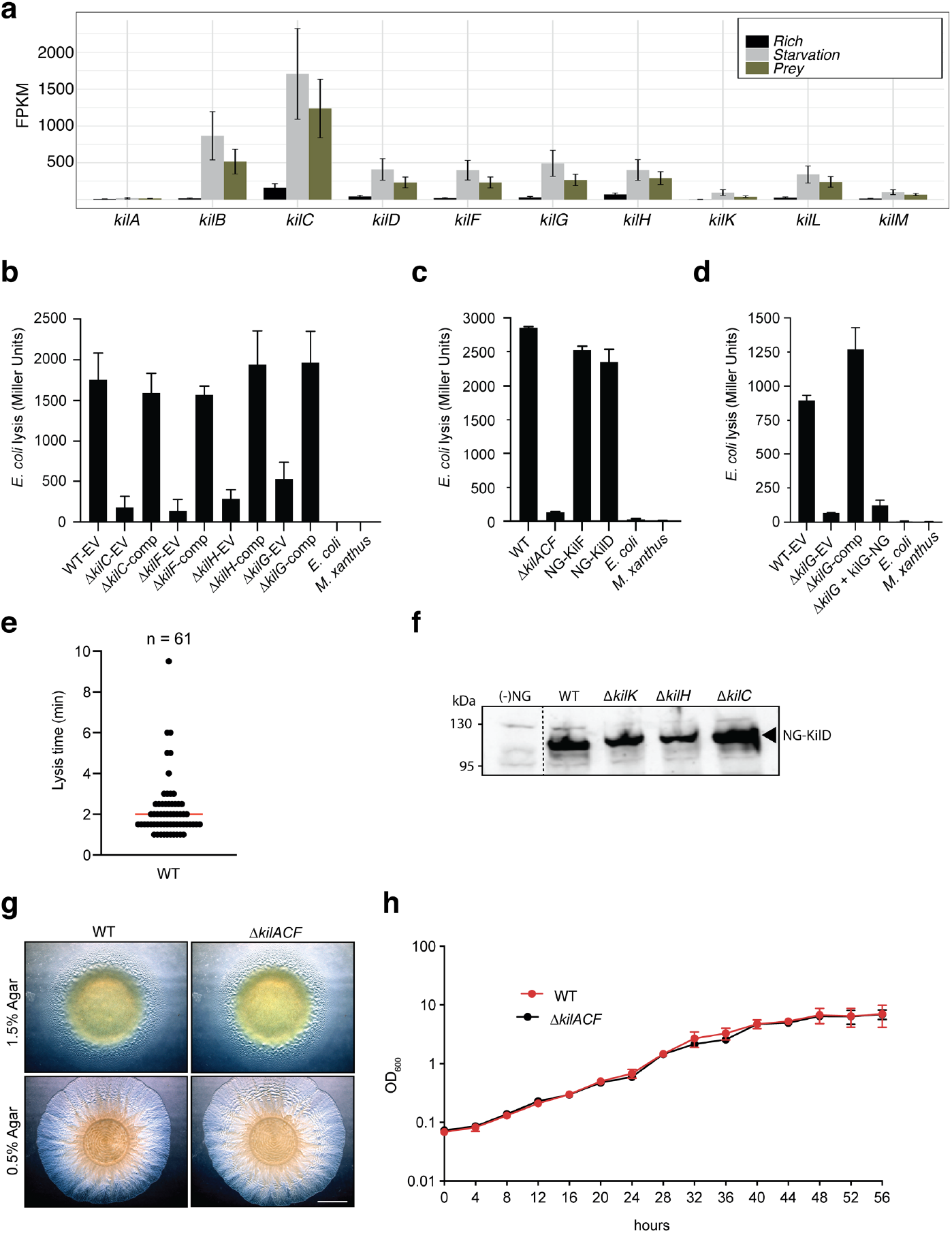
Functional analysis of the *kil* genes. **a:** The *kil* genes are expressed during starvation. RNA-seq analysis of *kil* gene expression in rich medium, starvation medium and starvation medium with live prey cells extracted and computed from data by Livingstone *et al*.^61^. For each gene and condition the data is compiled from three independent biological replicates. Note addition of prey does not change the expression profile which is significantly induced by starvation alone. **b:** CPRG colorimetric assay. In a 96-well plate, the different *kil* strains transformed with pSWU19-EV (Empty Vector) or complemented (comp) with a pSWU19 carrying the different *kil* genes were incubated with *E. coli* in liquid. After 24-hour incubation, β-galactosidase activities (expressed as Miller Units) of the different cell lysates were measured. The *kil* mutants ectopically expressing the different *kil* genes under the control of the *pilA* promoter presented a restored predation phenotype similar to WT-EV control. *E. coli* and *M. xanthus* alone were used as negative controls. This experiment was independently performed twice with at least two experimental replicates per strain each time. Error bars represent the standard deviation to the mean. **c:** the strains expressing neon Green (NG) fusions of KilD or KilF have predation phenotype similar to wild-type in a CPRG colorimetric assay. This experiment was independently performed twice. Error bars represent the standard deviation to the mean. **d:** Δ*kilG* strain expressing KilG-NG is defective in predation. CPRG colorimetric assay. In a 96-well plate, Δ*kilG* transformed with pSWU19-EV, pSWU19-*PpilA*-*kilG* or pSWU19-*PpilA*-*kilG-NG* gene was incubated with *E. coli* in liquid. After 24-hour incubation, β-galactosidase activities (expressed as Miller Units) of the different cell lysates were measured. Only Δ*kilG* pSWU19-*PpilA*-*kilG* complemented strain presented restored predation phenotype similar to the WT-EV control. *E. coli* and *M. xanthus* alone were used as negative controls. This experiment was performed once with four experimental replicates per strain. Error bars represent the standard deviation to the mean. **e:** Time to lysis after cluster formation. Time to lysis was determined by first monitoring cluster formation and then loss of contrast by the prey cell. The measurements were performed over two biological replicates. The median is shown as a red bar. **f:** Stable expression of NG-KilD in different mutant backgrounds. NG-KilD is detected at the expected molecular weight by the anti-neon Green antibody. (-) NG: DZ2 *Myxococcus* cell extracts that do not express neon Green. Dotted line indicates gel splicing. **g:** Growth and motility of WT and Δ*kilACF* mutant strains on agar supporting both A- and S-motility (1.5%) and S-motility only (0.5%). Scale bar = 2 mm. **h:** Growth curves of WT and Δ*kilACF* mutant in CYE rich medium. The measurements were performed over three biological replicates. Error bars represent the standard deviation to the mean.

**Figure S5:**
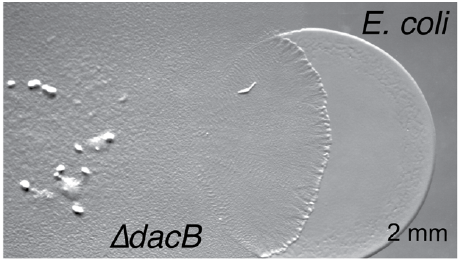
Predation phenotype of a *Myxococcus* D,D-decarboxylase mutant^42^. Colony plate assays showing invasion of an *E. coli* prey colony (dotted line) 48 hours after plating by a *dacB* mutant. Scale bar = 2 mm.

## Legends to Supplemental Movies

**Movie S1: Invasion of *E. coli* colonies by WT *Myxococcus* cells**. This movie was taken at the interface between the two colonies during invasion. The movie is a 8x compression of an original movie that was shot for 10 hours with a frame taken every 30s at 40x magnification. To facilitate *Myxococcus* cells tracking, the wild-type strain was labeled with the mCherry fluorescent protein.

**Movie S2: A-motility is required for prey invasion**. This movie was taken at the interface between the two colonies during invasion. The movie is a compression of an original movie that was shot for 10 hours with a frame taken every 30s at 40x magnification. To facilitate *Myxococcus* cells tracking, the A^-^S^+^ (*ΔaglQ*) strain was labeled with the mCherry fluorescent protein.

**Movie S3: Prey invasion by A-motile cells in “arrowhead” formations**. Focal adhesions and thus active A-motility complexes were detected with an AglZ-Neon green fusion. The movie contains 51 frames taken every 30 seconds at 100x magnification. Shown side-by-side are fluorescence images, fluorescence overlaid with phase contrast and MiSiC segmentation (lysing *E. coli* cells are colored magenta and blue).

**Movie S4: A *Myxococcus* cell kills an *E. coli* cell by contact**. The *Myxococcus* cell expresses AglZ-nG and the *E. coli* cell expresses mCherry. Shown side-by-side are fluorescence images and MiSiC segmentation (*Myxococcus*: green, *E. coli*: magenta). The movie contains 20 frames taken every 30 seconds at 100x magnification.

**Movie S5: NG-KilF cluster formation in contact with *E. coli* prey cells**. Shown is an overlay of the fluorescence and phase contrast images of a motile *Myxococcus* cell in predatory contact with three *E. coli* cells. The movie was shot at 100x magnification objective for 30 minutes. Pictures were taken every 30 seconds.

**Movie S6: KilG-NG cluster formation in contact with *E. coli* prey cells**. Shown is an overlay of the fluorescence and phase contrast images of a motile *Myxococcus* cell in predatory contact with three *E. coli* cells. The movie was shot at 100x magnification objective for 9 minutes. Pictures were taken every 30 seconds.

**Movie S7: NG-KilD cluster formation in contact with *E. coli* prey cells**. Shown is an overlay of the fluorescence and phase contrast images of a motile *Myxococcus* cell in predatory contact with three *E. coli* cells. The movie was shot at 100x magnification objective for 15 minutes. Pictures were taken every 30 seconds.

**Movie S8: NG-KilD clusters form in a *kilC* mutant but no motility pauses and prey cell lysis can be observed**. Shown is an overlay of the fluorescence and phase contrast images of a motile *Myxococcus* cell in predatory contact with three *E. coli* cells. The movie was shot at 100x magnification objective for 8 minutes. Pictures were taken every 30 seconds.

**Movie S9: a Δ*kilACF* still invades but does not kill *E. coli* prey cells**. This movie was taken at the interface between the two colonies during invasion. The movie is a 4x compression of an original movie that was shot for 4.5 hours with a frame taken every 30s at 40x magnification. To facilitate *Myxococcus* Δ*kilACF* cells are labeled with the mCherry fluorescent protein.

**Movie S10: Predatory cells division and tracking during invasion of prey colony**. To follow cell growth and division at the single cell level during prey invasion, WT cells were mixed with a WT strain expressing the mCherry at a 50:1 ratio and imaged every 30 seconds at 40x magnification for up to 10 hours within non-labeled prey colonies. Cell growth was measured by fitting cell contours to medial axis model followed by tracking under Microbe-J. Real time of the track for the example cell: 95 min.

**Movie S11: NG-KilD cluster formation in contact with *Caulobacter crescentus* prey cells**. Shown is an overlay of the fluorescence and phase contrast images of a motile *Myxococcus* cell in predatory contact with a *C. crescentus* cell. The movie was shot at 100x magnification objective for 7 minutes. Pictures were taken every 30 seconds.

**Movie S12: NG-KilD cluster formation in contact with *Salmonella typhimurium* prey cells**. Shown is an overlay of the fluorescence and phase contrast images of a motile *Myxococcus* cell in predatory contact with an *S. enterica* Typhimurium cell. The movie was shot at 100x magnification objective for 20 minutes. Pictures were taken every 30 seconds.

**Movie S13: NG-KilD cluster formation in contact with *Bacillus subtilis* prey cells**. Shown is an overlay of the fluorescence and phase contrast images of a motile *Myxococcus* cell in predatory contact with a *B. subtilis* cell. The movie was shot at 100x magnification objective for 30 minutes. Pictures were taken every 30 seconds.

**Movie S14: *Pseudomonas aeruginosa* is not lysed by *Myxococcus* and does not induce NG-KilD cluster formation**. Shown is an overlay of the fluorescence and phase contrast images of a motile *Myxococcus* cells mixed with *Pseudomonas* cells. The movie was shot at 100x magnification objective for 30 minutes. Pictures were taken every 30 seconds.

